# Large-scale analyses of human microbiomes reveal thousands of small, novel genes and their predicted functions

**DOI:** 10.1101/494179

**Authors:** Hila Sberro, Nicholas Greenfield, Georgios Pavlopoulos, Nikos Kyrpides, Ami S. Bhatt

## Abstract

Small proteins likely abound in prokaryotes, and may mediate much of the communication that occurs between organisms within a microbiome and their host. Unfortunately, small proteins are traditionally overlooked in biology, in part due to the computational and experimental difficulties in detecting them. To systematically identify novel small proteins, we carried out a large comparative genomics study on 1,773 HMP human-associated metagenomes from four different body sites (mouth, gut, skin and vagina). We describe more than four thousand conserved protein families, the majority of which are novel; ~30% of these protein families are predicted to be secreted or transmembrane. Over 90% of the small protein families have no known domain, and almost half are not represented in reference genomes, emphasizing the incompleteness of knowledge in this space. Our analysis exposes putative novel ‘housekeeping’ small protein families, including a potential novel ribosomally associated protein, as well as ‘mammalian-specific’ or ‘human-specific’ protein families. By analyzing the genomic neighborhood of small genes, we pinpoint a subset of families that are potentially associated with defense against bacteriophage. Finally, we identify families that may be subject to horizontal transfer and are thus potentially involved in adaptation of bacteria to the changing human environment. Our study suggest that small proteins are highly abundant and that those of the human microbiome, in particular, may perform diverse functions that have not been previously reported.

## Introduction

Numerous studies have correlated the microbiota with human health and disease states (Ley et al., 2006; Liang et al., 2018; Manrique et al., 2017; Reid, 2018; Sekirov et al., 2010; Shreiner et al., 2015; Verma et al., 2018; Weyrich et al., 2015). In most studies an amplicon-based approach is used, often leveraging sequencing of the 16S ribosomal RNA gene, to characterize the taxonomic composition of a sample. However, human microbiome research is transitioning from a primarily descriptive science to a more mechanistic science (Gilbert et al., 2018). To support this transition, there is an ongoing shift to whole-metagenome shotgun (WGS) sequencing projects, such as the National Institutes of Health Human Microbiome Project (HMP) (Lloyd-Price et al., 2017). In WGS studies DNA is extracted, sheared and sequenced, facilitating the identification of genes encoded by the underlying microbial community (Chen et al., 2010; Consortium, 2010; Li et al., 2014; Lloyd-Price et al., 2017; Qin et al., 2010; Ranjan et al., 2016). While accumulating WGS studies have illuminated the remarkable genetic diversity encoded by human-associated microbes, our ability to link specific genes, molecules or pathways to phenotypes is still lagging behind (Koppel and Balskus, 2016). One of the major challenges in linking genes to phenotypes is that the process of gene annotation is incomplete, and overlooks an entire class of potentially important genes.

Small open reading frames (sORFs) and the small proteins they encode, here defined as proteins of ≤50 amino acids in length, have traditionally been ignored (Duval and Cossart, 2017; Storz et al., 2014; Su et al., 2013). In the classical process of genome annotation, ORFs are predicted by relying on attributes of the sequence (e.g. presence of start and stop codons) or through homology to other proteins. However, sORFs prediction is bioinformatically challenging as it is difficult to distinguish true protein coding ORFs from the numerous random in-frame genome fragments. To distinguish genuine protein-coding ORFs from randomly occurring fragments, most prediction tools require a minimum ORF length, which results in databases that almost entirely lack sORFs. The computational challenges associated with the identification of sORFs are further augmented in the prediction of their putative function. In mutational screens, where the likelihood of mutating a gene is proportional to gene length, it is proportionally less likely that mutagens will target a sORF and allow for characterization of the loss-of-function effects. Finally, classical biochemical approaches are usually not optimized to detect small proteins and experiments that rely on databases, such as mass spectrometry, will fail to identify small proteins if their sequences are not present in reference databases (Duval and Cossart, 2017; Storz et al., 2014; Su et al., 2013).

Despite the systematic bias against the discovery and characterization of proteins encoded by sORFs, recent studies in both eukaryotes and prokaryotes have elucidated interesting functions for this class of proteins (reviewed in Couso and Patraquim, 2017; Duval and Cossart, 2017; Kemp and Cymer, 2014; Storz et al., 2014). In eukaryotes, small proteins play roles in development (Chng et al., 2013; Galindo et al., 2007; Kondo et al., 2010), DNA repair (Slavoff et al., 2014), calcium homeostasis (Anderson et al., 2015, 2016; Magny et al., 2013; Nelson et al., 2016), myoblast fusion (Bi et al., 2017), metabolism (Lee et al., 2015) and mRNA turnover (D’Lima et al., 2017). In prokaryotes, small proteins play roles as ‘housekeeping’ ribosomal proteins (Yutin et al., 2012), chaperones (Martin et al., 2015; Smaldone et al., 2012) and cell division proteins (Handler et al., 2008; Hobbs et al., 2010; Modell et al., 2011), and in signal transduction (Lippa and Goulian, 2009; Rowland et al., 2004), transport (Gaßel et al., 1999; Hobbs et al., 2012; Lloyd et al., 2017), regulation of membrane-bound enzymes (Choi et al., 2012; Kato et al., 2012; Sun et al., 2012) and bacterial stress response (Impens et al., 2017; Martin et al., 2015). Small microbial proteins also take part in communication and warfare between cells. For example, quorum sensing, a communication mechanism that influences sporulation, competence, antibiotic production, biofilm formation and virulence is mediated by small proteins (Moreno-Gámez et al., 2017). In addition, some of the antimicrobial peptides produced by bacteria against other microorganisms are small proteins (Cotter et al., 2013). Finally, recent work has revealed a small bacteriophage protein that influences whether phages replicate and lyse the host or lysogenize and remain dormant (Erez et al., 2017).

The HMP datasets represent a rich resource for sORF discovery, and we thus sought to enumerate and classify the small proteins encoded by the healthy human microbiome. We were intrigued by this class of typically overlooked proteins since they can be readily synthesized and may represent useful model systems for protein folding simulation given that they generally consist of a single small domain (Polticelli et al., 2001). Finally, these proteins are of interest in the pharmaceutical field as potential drugs (Anderson et al., 2015; Fosgerau and Hoffmann, 2015; Lau and Dunn, 2018).

To identify novel sORFs with high precision and minimize false positive findings, we leveraged the concept that sORFs that are transcribed, translated and functional likely have protein sequences that are conserved across species. Our analysis reveals 4,539 small protein families encoded by human-associated microbes. Based on the rigorous method used for sORF identification, we have high confidence that this database of approximately four thousand small proteins does not contain spurious predictions. Intriguingly, very few proteins found to be abundant in human-associated microbes have been thoroughly characterized.

For each family of homologous small proteins, we provide detailed information about its taxonomic classification, prevalence across human-metagenomes and across body sites, predicted cellular localization (secreted/transmembrane) and the presence or absence of homologs of the families among ~6,000 non-human metagenomes. Since gene context can inform predictions of function in bacterial genomes, we also provide information about genes that are encoded in the vicinity of representatives of each sORF family.

Using this integrated approach, we highlight several novel small proteins with diverse predicted functions. We describe a group of small proteins that are extremely prevalent, one of which is likely to be a novel ribosomally associated protein and another that is an abundant post-translationally modified small protein; we expose small proteins that are potentially involved in the crosstalk with the host or with other microbial cells; we find small proteins that are likely subject to horizontal gene transfer and highlight small protein families that could be related to defense against phage or against other bacteria; finally, we enumerate small protein families originating from the non-bacterial fraction of the human microbiome.

Altogether, we present a rich resource of sORFs encoded by human-associated microorganisms, and provide insights into the potential functions of these proteins, which we anticipate will inform future studies in the human associated proteome. Building an accurate understanding of the full coding potential encoded by the human microbiome, including the thus far overlooked sORFs, is a fundamental step towards understanding of the mechanisms that underlie the role of the microbiome in health and disease.

## Results

### Only a small subset of well characterized small proteins are relevant to the human microbiome

To explore the small proteins encoded by the human microbiome, we first asked whether small proteins that have already been studied and characterized are present in the human microbiome. Small proteins that have been studied in depth generally originate from model organisms (reviewed in Duval and Cossart, 2017; Storz et al., 2014). To infer their potential relevance to the human microbiome, we sought to identify those sORFs that are also encoded by organisms that are prevalent in human-associated microbial communities. In order not to limit our search to species that have a reference genome, we undertook a reference-free approach and conducted our analysis directly on metagenomic sequencing data. We chose to work with the recently released HMPI-II dataset (Lloyd-Price et al., 2017) that includes shotgun sequenced human-associated metagenomes from healthy subjects. We used the gene prediction tool MetaProdigal (Hyatt et al., 2012), that is suitable for detection of genes in short coding sequences of unknown origin, to annotate all open reading frames (ORFs) on 128,368,337 contigs spanning more than 180 billion base pairs (bps) of sequenced DNA from 1,773 metagenomes from 263 individuals (Supplementary Table S1) sampled from four different major body sites (mouth, gut, skin and vagina, Supplementary Table S2). We modified MetaProdigal’s default parameters to identify ORFs as short as 15 bp. From the resulting set of ORFs, we filtered out ORFs that encode for proteins that are greater than 50 amino acids in length, without any lower limit threshold, resulting in a set of 2,514,099 small ORFs ranging in size from eight amino acids to 50 amino acids (Figure 1A).

**Figure 1.**
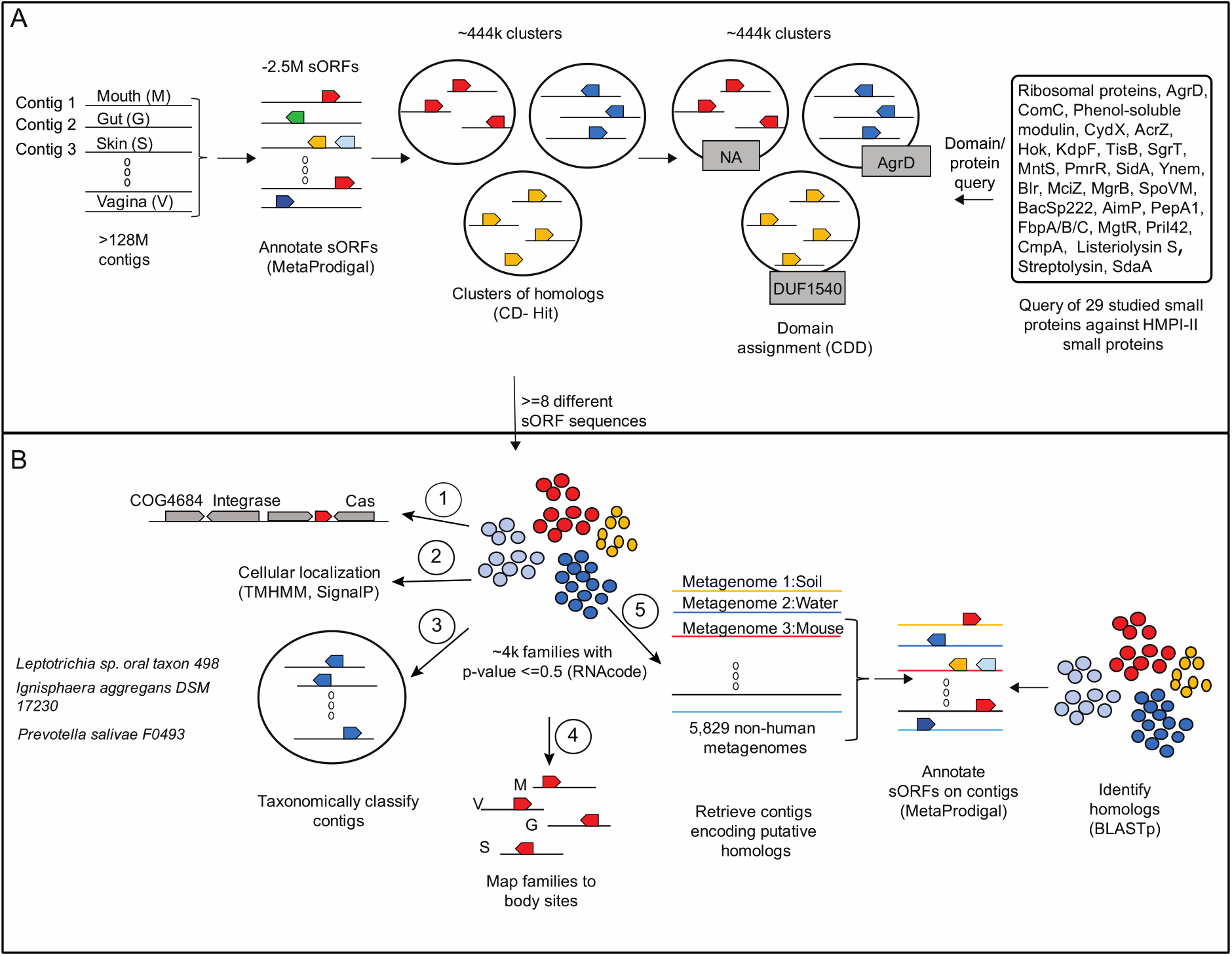
Small protein discovery and characterization pipeline applied to HMPI-II metagenomic data. **A**. Identification of 29 known small proteins in HMPI-II metagenomes. More than 128M contigs were annotated using MetaProdigal with a lower size limit of five amino acids. The small proteins were then clustered using CD-Hit based on amino acid similarity and protein length. Representatives of each of the ~444 thousand clusters were queried against the Conserved Domain Database (CDD), to assign domains to clusters. The list of CDD domains was then queried for the small known proteins that have an assigned domain. Known small proteins that do not have an assigned domain or that failed the domain search were queried against HMPI-II small proteins using BLASTp. **B**. Identification and characterization of HMPI-II small proteins. RNAcode was used to assign p-values to the ~444 thousand clusters. The following analyses were conducted on the ~4k protein families whose p-value was ≤0.05: 1) identification of neighboring genes on longest contig associated with each family 2) prediction of signal peptide and transmembrane domains 3) taxonomic classification of contigs encoding each of the small protein families 4) assignment of small protein families to body sites 5) identification of homologs of small protein families in non-human metagenomes.

We queried a set of 29 known small proteins that have been studied in depth (reviewed by Duval and Cossart, 2017; Storz et al., 2014) (Table 1) as well as a set of small ribosomal proteins, and attempted to identify homologs of these known small proteins among the ~2.5M putative small proteins predicted by our sORF annotation with MetaProdigal. Whenever possible we used a domain-based approach (conserved domain search using RPS-BLAST) that would detect even distant homologs (Altschul et al., 1997) and we used a sequence-based approach (BLASTp) for small known proteins that have not been assigned a protein domain. Only 12 of the 29 small proteins have an assigned a protein domain (AcrZ, CydX, KdpF, AgrD, ComC, MciZ, MgrB, SpoVM, SgrT, Hok, TisB, phenol-soluble modulins as well as small ribosomal proteins). To determine which of the ~2.5M putative small proteins contains one of these domains, we first had to assign domains to the putative HMP small proteins. To reduce computational load associated with analysis of such large amounts of sequences, we started by clustering all ~2.5M putative small protein sequences based on sequence and length similarity using CD-Hit (Fu et al., 2012), a tool which takes a sequence database as input and produces a set of ‘non-redundant’ representatives as well as a set of sequence families associated with each representative. This resulted in 444,054 clusters of homologous sequences. We then queried each of the 444,054 families’ representatives against the Conserved Domain Database (CDD), which serves as an umbrella to multiple datasets including COG (Clusters of Orthologous Groups of Proteins), Pfam (Protein Families), SMART (Simple Modular Architecture Research Tool) and TIGRFAM (The Institute of Genomic Research’s database of protein families) (Marchler-Bauer et al., 2011, 2017) (Figure 1A and Supplementary Table S3).

**Table 1.**
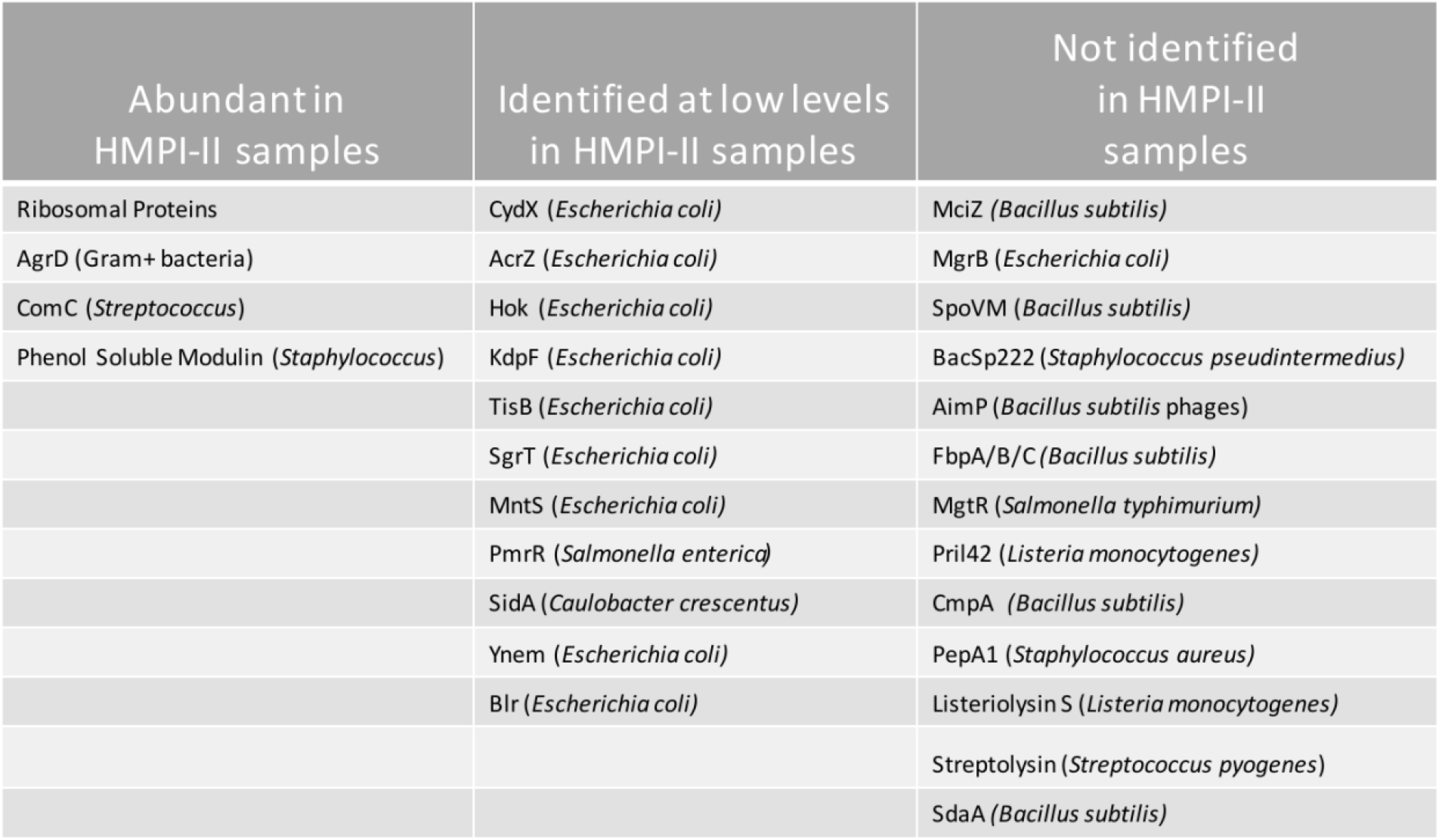
Representation of known small proteins in HMPI-II data. Known proteins were queried against CDD-assigned domains of all 444,054 representatives whenever they have an assigned domain and against all protein sequences of the ~444,054 representatives in case the domain search failed and when known protein is not assigned a known domain. Names of organisms in parentheses indicate the model organism in which small protein was mainly studied.

| Abundant in HMPI-II samples | Identified at low levels in HMPI-II samples | Not identified in HMPI-II samples |
| --- | --- | --- |
| Ribosomal Proteins | CydX (*Escherichia coli*) | MciZ (*Bacillus subtilis*) |
| AgrD (Gram+ bacteria) | AcrZ (*Escherichia coli*) | MgrB (*Escherichia coli*) |
| ComC (*Streptococcus*) | Hok (*Escherichia coli*) | SpoVM (*Bacillus subtilis*) |
| Phenol Soluble Modulin (*Staphylococcus*) | KdpF (*Escherichia coli*) | BacSp222 (*Staphylococcus pseudintermedius*) |
|  | TisB (*Escherichia coli*) | AimP (*Bacillus subtilis* phages) |
|  | SgrT (*Escherichia coli*) | FbpA/B/C (*Bacillus subtilis*) |
|  | MntS (*Escherichia coli*) | MgtR (*Salmonella typhimurium*) |
|  | PmrR (*Salmonella enterica*) | Pril42 (*Listeria monocytogenes*) |
|  | SidA (*Caulobacter crescentus*) | CmpA (*Bacillus subtilis*) |
|  | Ynem (*Escherichia coli*) | PepA1 (*Staphylococcus aureus*) |
|  | Blr (*Escherichia coli*) | Listeriolysin S (*Listeria monocytogenes*) |
|  |  | Streptolysin (*Streptococcus pyogenes*) |
|  |  | SdaA (*Bacillus subtilis*) |

Only ~4.5% (113,693/2,514,099) of the putative small proteins, spanning ~0.5% (2,225/444,054) of the clusters, could be assigned a known domain (Supplementary Table S3). Among the assigned sequences, the most common type of domains that could be identified are of diverse small ribosomal proteins, assigned to ~64% of all domain-assigned small proteins (72,982/113,693). In addition, we found small proteins that were identified in model organisms that are known inhabitants of the human microbiome. For example, ~3.5% of small proteins that were assigned a domain (3,930/113,693) were homologous to the extensively studied quorum sensing small protein, *Staphylococcal* AgrD. Homologs of AgrD were clustered into 153 clusters, suggesting rapid evolution of the proteins since the sequences were divergent enough to cluster independently of one another. This is in line with what has been previously documented for AgrD; specifically, it has been shown that AgrD has a highly variable sequence, which is presumed to result in varied signaling specificities (Hyatt et al., 2012).

For the known small proteins that do not have an assigned domain and for those that failed the domain search, we used BLASTp (Camacho et al., 2009) to search for their sequence against all representatives. Additional information about representation of known small proteins in our dataset can be found in the Supplementary Table S4 and in Table 1.

Overall, our analysis of well-studied small proteins suggests that there is a small overlap between small proteins that have been extensively characterized in model organisms and those that are abundant and potentially important in the human microbiome. Well studied proteins that were abundant in our dataset (such as AgrD and ComC) are encoded by commonly studied organisms that are often constituents of the healthy microbiome (such as *Staphylococcus* and *Streptococcus*, respectively), making it unsurprising that we identified them in our human-associated microbiome data set.

### The ~4k small protein families of the human microbiome are in most part unknown

We were intrigued that such a small proportion of previously described small proteins were present in the human-associated microbiomes. We thus sought to better understand what types of small proteins exist in this unexplored space.

As a first step, we revisited the 444,054 clusters (Table S3) of potential small proteins that were generated in the previous step of our analysis (Figure 1A). Since most were not assigned a known functional domain, which raised the concern for potential presence of spurious sORFs in this dataset, we sought to enrich for families that are more likely to be genuine, protein-coding families.

To enrich for protein-coding sequences, we used RNAcode (Washietl et al., 2011), a gene predictor program that distinguished between coding and non-coding sequences by evaluating evolutionary signatures, including synonymous/nonsynonymous mutations within multiple sequence alignments of putative homologs. This program was shown to perform substantially better and reach almost perfect discrimination between coding and non-coding sequences when the number of input sequences is ≥8 (Washietl et al., 2011). Thus, we applied RNAcode on the 11,715 clusters that contained ≥ 8 different DNA sequences (putative homologs). Using a p-value threshold of ≤0.05, we identified 4,539 clusters from the 11,715 (contain 467,538 small protein sequences) that are predicted to be bona fide sORFs (Figure 1A, Supplementary Table S5). We denote these clusters ‘small protein families’ and these ‘small protein families’ are subjected to further downstream analyses hereafter. Supplementary figures 3 and 4 display additional attributes of the families.

The ~4k families that were identified using RNAcode are a subset of the 444,054 clusters we started our analysis with, hence have already been assigned a protein domain, whenever possible. Reassuringly, the ~4k family subset is significantly enriched for small protein families that were assigned a protein domain in our previous analysis (p < 1×10^-5^ Fisher Exact Test): among the 4,539 small protein families, 4% (192/4,539) were assigned a domain (compared to 0.5% of the 444,054 clusters), (Figure 2A-B). These families contain 12% of the 467,538 small proteins (compared to 4.5% of the 2,514,099 in the initial database). The enrichment for likely true positive sORFs with the application of the RNAcode algorithm supports its inclusion in our approach to identify genuine small protein families. Still, ~96% (4,351/4,539) of small protein families were not assigned a CDD domain (Figure 2A, Supplementary Table S5), some of which are actually encoded by a large number of species (Figure 2C, Supplementary Table S5), emphasizing the incompleteness of knowledge in small protein domains space.

**Figure 2.**
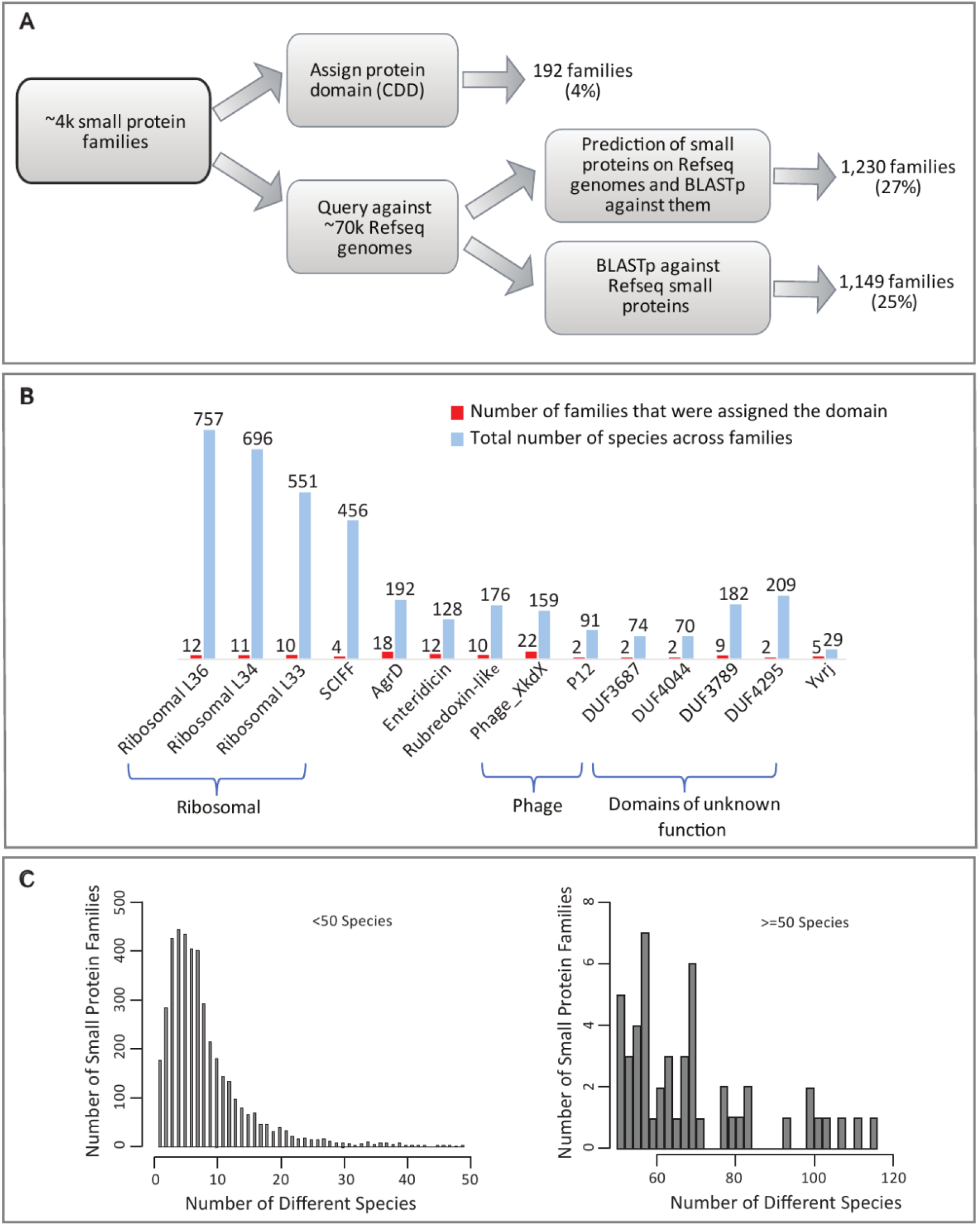
Most of the ~4k families, some of which are very abundant, are not assigned a known protein domain nor are they represented in RefSeq genomes. **A**. Pipeline to identify families that do not have an assigned domain and families that are not represented in RefSeq genomes. Upper part: only a small subset of the ~4k small protein families were assigned a protein domain (identified by RPS-blast against CDD position specific scoring matrices, PSSMs). Lower part: Representatives of all ~4k families were blasted against ~3M small RefSeq annotated proteins originating from ~70k RefSeq genomes and against ~7M putative small proteins that we annotated using Prodigal with adjusted thresholds. The second step allowed the identification of an additional set of homologs that are encoded but not annotated in RefSeq genomes. **B**. Domains identified among ~4k families. Domains that were classified to ≥5 families and/or ≥50 species are shown. A complete list of domains can be found in Supplementary Table S5. **C**. Number of species encoding small proteins of families with no known domain are shown in histogram.

In addition to identifying which of the well characterized small proteins are represented in the human microbiome, we also asked an orthogonal question: what proportion of the ‘high confidence’ sORF families are found in reference genome databases such as RefSeq (Pruitt et al., 2007)? To answer this question, we performed sequence similarity searches of all 4,539 representative proteins in the ~4k families against proteins of ≤50 amino acids annotated in 69,681 RefSeq bacterial reference genomes. Our search showed that only ~25% of the small protein families (1,149/4,539) in our dataset are annotated in RefSeq genomes (Figure 2A and Supplementary Table S5).

Since sORFs are often excluded in annotation process, we postulated that at least some of the small proteins in our data set do have homologs that were not annotated in RefSeq. We therefore re-annotated all 69,681 RefSeq genomes with a permissive size threshold to include all potential sORFs. Indeed, this step revealed an additional set of 1,230 (~27%) small protein families that do have RefSeq homologs that were previously not annotated (Figure 2A and Supplementary Table S5). Still, for 48% (2,164/4,539) of the small protein families, we could not identify any homologs (annotated or not) among RefSeq genomes (Figure 2A). This gap confirms that any effort to comprehensively identify candidate novel small proteins of the human microbiome would be very limited if applied only to genomes from reference databases that have, generally speaking, a limited representation of human-associated microbes.

Altogether, our analysis suggests that the vast majority of small protein families reported here are unknown: they are not well represented among reference genomes and most do not have an assigned protein domain.

Since most families in our dataset do not have an assigned protein domain nor do they have well characterized homologs from which we can try to infer function, we used the following approaches to provide insight into the potential functions of these small proteins (Figure 1B). Briefly, we analyze the taxonomy, the prevalence of families across samples, body sites, and non-human metagenomes, we predict their cellular localization (secreted/transmembrane/other), and since gene context is very informative in deciphering function in bacterial genomics, we also provide information about genes that are encoded in close vicinity (up to 10 genes away) to the small proteins. In the next sections we focus on specific classes of small proteins. The examples we provide are probably merely the ‘tip of the iceberg’ of potential functions encoded by small proteins. Supplementary Table S5 provides detailed information about all families.

### Putative novel ‘housekeeping’ small protein families among human-associated microbes

We next sought to identify small protein families that could be playing housekeeping roles. We posited that such protein families would be highly conserved across the tree of life, and thus searched for small protein families that are prevalent across species. To characterize the taxonomic distribution of families, we classified each of the contigs that encoded small proteins against a set of 83,701 microbial reference genomes, consisting of 53,193 Bacteria, 27,020 Viruses, 1,892 Eukaryota (fungi and protozoa) and 1,756 Archaea genomes, using the k-mer based One Codex platform (Minot et al., 2015). Briefly, every k-mer in each contig was classified to its lowest common ancestor (LCA) across the One Codex database. Each contig was then assigned based on the results of its constituent k-mers. This contig-level classification was made by determining the taxonomic ID with the highest weighted root-to-leaf path across all k-mers within the contig.

Given that housekeeping proteins are expected to be conserved across the tree of life, we focused our next analysis on the 14 most prevalent families, that are encoded by ≥100 species (Supplementary Table S5 and Figure 3A). First, we characterized the taxonomic composition of these families. In all 14 small protein families, the average percentage of k-mers that could be classified is >10%, implying that classification is likely reliable in these families. Whereas most families in the overall dataset are taxonomically unique to one (2,353, 52%) or two (1,183, 26%) phyla, there is a strong enrichment among the 14 most prevalent families for presence in multiple phyla (Figure 3B), suggesting a role that is not clade-specific. Second, we determined whether these families are specific to a particular ecological niche. To characterize the niches in which small proteins are encoded, we mapped each family to the body sites in which homologs of the family were identified. Whereas most small protein families are identified uniquely in mouth (1,188, 26%) or gut (2,220, 48%) (Supplementary Table S5), thirteen of the 14 most prevalent families were identified in ≥3 body sites, suggesting a role that is not niche specific (Figure 3A). Since the HMP data resource we used for this study has a relatively limited representation of skin and vagina samples (Supplementary Table S2), it is possible that families that seem absent from one of these body sites are actually present but not detected.

**Figure 3.**
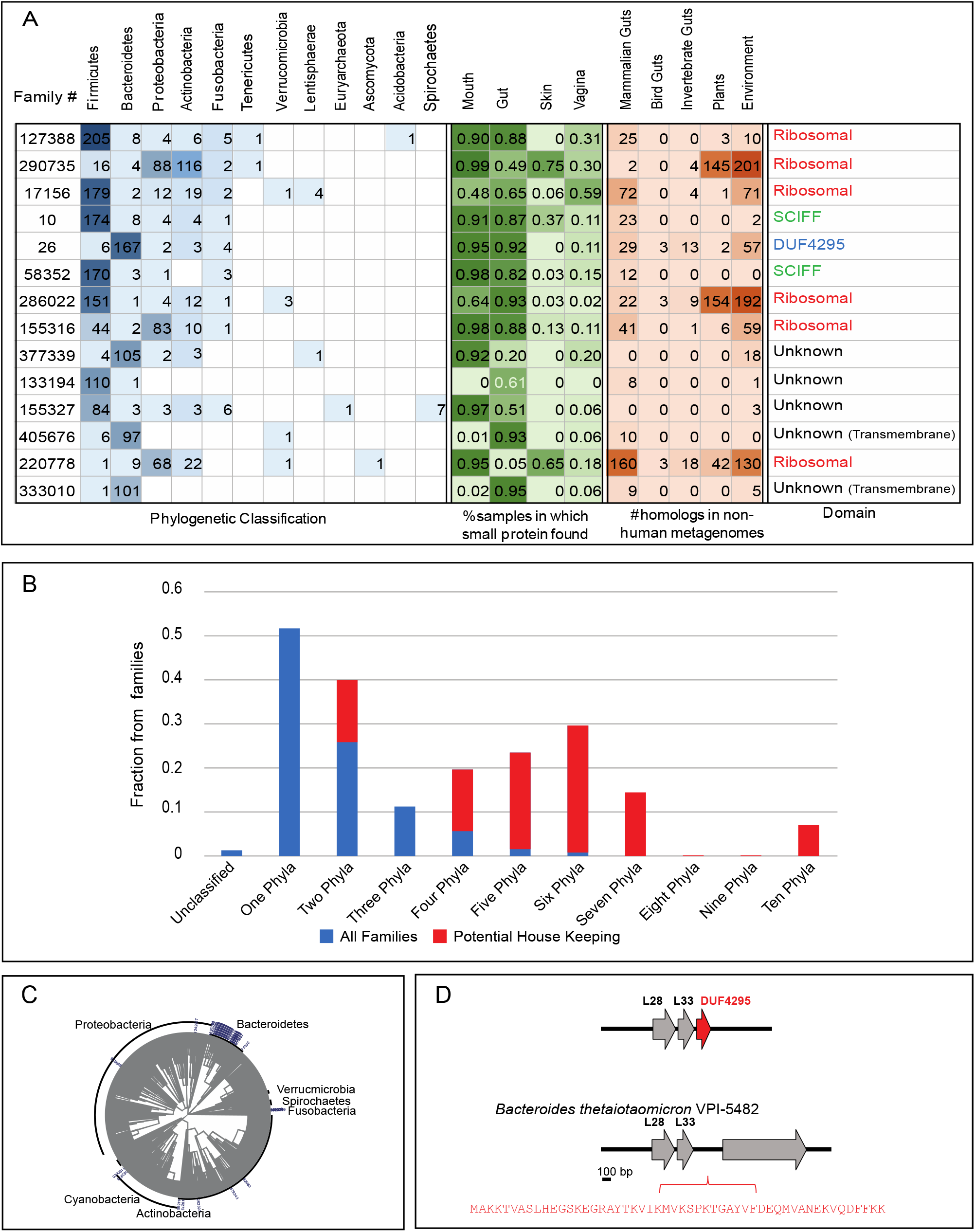
Small protein families that stand out in their taxonomic prevalence. **A**. Most abundant families. Each row represents one of the 14 families that were identified in ≥100 species. The taxonomic distribution of the 14 families is presented in the blue table, the prevalence among body sites is presented in the green table and the number of homologs identified in non-human metagenomes is presented in the brown table. Potential novel ribosomal is family #26. When multiple homologs were mapped to the same taxa, it is counted as one event in this table. **B**. The fraction of families assigned to different number of phyla for the 14 potential housekeeping (red) and the 4,525 remaining families (blue) is shown. For example, >50% of the non-housing-keeping families were assigned to one phyla vs zero housekeeping families that were assigned to one phylum. **C-D**. Potential novel ribosomal protein. **C**. Phylogenetic tree of family #26. **D**. The genomic neighborhood of DUF4295 (family #26) next to two know ribosomal proteins is illustrated. In *Bacteroides thetaiotaomicron (B. theta)* VPI-5482 it is encoded in the intergenic region downstream of these genes (locus tags BT0914 and BT0915).

Finally, positing that true housekeeping genes are likely to be conserved among a broad range of ecological niches, we tested whether these 14 prevalent families are more likely to have homologs in non-human metagenomes than the remainder of the ~4k small protein families defined earlier. To do so, we checked for sequence homology of the ~4k small proteins within a set of 5,829 non-human metagenomes, including mammalian and bird gut metagenomes, as well as environmental samples of different types (Methods and Supplementary Table S6). While we could not identify homologs in non-human metagenomes for the majority of small protein families (3,551, 78%), we were able to identify homologs in at least one non-human environment for all 14 families (Figure 3A). Altogether, the taxonomic abundance and the existence in multiple niches of these 14 families suggest a role that is not clade or niche specific. These two traits are associated with proteins that play housekeeping roles. Indeed, among these 14 potential ‘housekeeping’ families, six encode for different ribosomal proteins, a known class of housekeeping proteins. Among the remaining eight families, three were assigned a CDD domain and five were not. Two of the CDD-assigned families (#10 and #58352) were assigned the ‘SCIFF’ (“six cysteines in forty-five residues”) domain. This domain is associated with a small ribosomally-synthesized natural product that is post translationally modified by a radical SAM maturase enzyme (Haft and Basu, 2011; Haft and Haft, 2017). The biological function of this small protein is unknown. Another domain, assigned to family #26, is a DUF4295 domain, which we address below. Finally, there are five families that were not assigned a protein domain, two of which are predicted to be transmembrane.

### Identification of a putative novel ribosomally associated protein prevalent among human associated microbes

Family #26 is among the 14 families that are very abundant that were assigned a domain of unknown function (Figure 3C-D). This family encodes a 50-amino acid protein and was assigned the domain DUF4295. It was detected in 182 species originating from four different phyla (Supplementary Table S5). We identified homologs of this protein in diverse non-human metagenomes and in a high percentage of gut and mouth samples, as well as in vaginal samples (Supplementary Table S5). DUF4295 drew our specific attention because the sORF is located in a strongly conserved genomic locus, downstream of two known ribosomal proteins, L28 and L33 (Figure 3D). In light of its wide phylogenetic distribution and genomic localization, we hypothesize that this small protein family is a novel small ribosomally associated protein that has thus far escaped detection. In the lab strain *Bacteroides thetaiotaomicron* VPI-5482, the small gene encoding this protein was not annotated, as is the case for many small proteins, but nevertheless is encoded in the intergenic region downstream these two genes (Figure 3D).

Interestingly, this family is likely larger than is represented by family #26, alone. We identified another family, family #7858, which also encodes for the DUF4295 domain (Supplementary Table S5). It is encoded by three different phyla but since these originate from a relatively small number of species (26 species), it did not pass the required threshold (requiring ≥100 species) described above. However, it displays significant sequence homology to family 26 (Supplementary Figure 5) and is also found next to two ribosomal proteins; it is found in 85% of mouth samples (but not in any gut samples) as well as in diverse non-human environments. Altogether, we hypothesize that DUF4295 is a novel ribosomally associated domain.

### Small protein families that are ‘core’ to specific body site or sites

Next, we asked which small protein families in our dataset could be playing roles that are associated with a specific body niche. To identify the body site(s) with which each family is associated, we mapped all contigs associated with the ~4 thousand protein families back to body site from which these contigs were assembled. Contigs originating from human gut samples are dominated by *Bacteroidaceae* and *Ruminococcacea* families; those from mouth samples are dominated by the *Streptococcus*, skin sORFs are dominated *Staphylococcus* and finally, vaginal sORFs are dominated by *Lactobacillus* and *Prevotella* (Supplementary Table S7). This is in line with previous studies that showed dominance of these taxonomic clades in these body sites (Lloyd-Price et al., 2016; Si et al., 2017), providing support to our classification results.

In terms of prevalence across samples, a total of 458 families (10%, 458/4,539) were identified in ≥ 50% of samples of at least one body site. Here we denoted these families as ‘core’ to that body site (Supplementary Table S5). In line with the fact that 68% of the sampled analyzed here are mouth samples, followed by 26% samples that are gut samples, mouth samples ‘contribute’ the largest amount of ‘core’ families, followed by mouth, skin and vagina (Figure 4). In most cases, “coreness” of a family is associated with a specific body site. It should be noted, that there are families that were detected in >50% of samples originating from one body site and in <50% of samples originating from another body site. In other words, families that are not ‘core’ to a specific body site are not necessarily completely absent from it (Supplementary Table S5).

**Figure 4.**
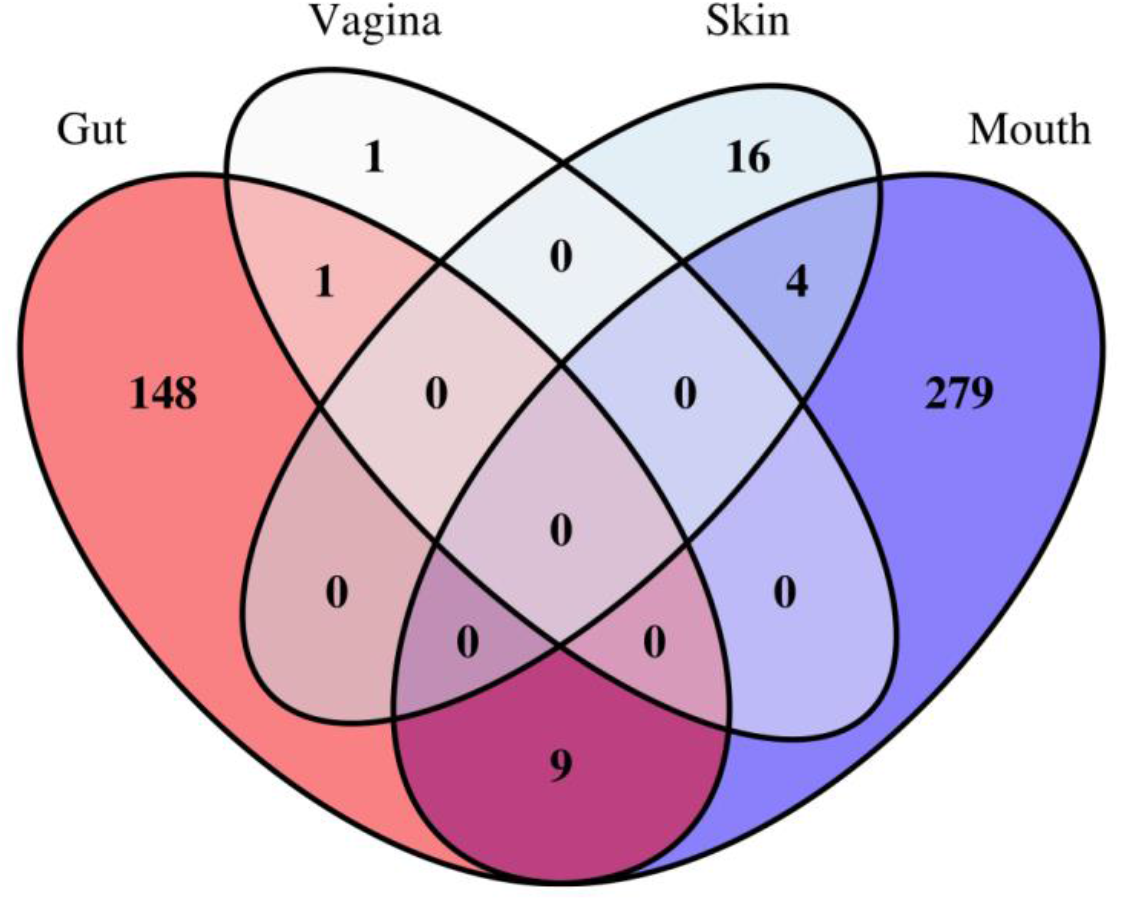
“Core” families across body sites. Venn-diagram representing the distribution of the 458 ‘core’ (identified in ≥ 50% from total samples of specific body site) small protein families across different body sites. Whereas mouth and gut share multiple small protein families that are found in ≥50% of the samples in both body sites, other pairs (e.g. skin ang gut) do not share core small genes, or share a very small number.

Only nine families are ‘core’ to both gut and mouth (Figure 4 and Supplementary Table S5). Not surprisingly, most (8/9) of these families are part of the list of potential housekeeping families (see above), that were identified based on their wide phyletic occurrence (i.e., in ≥100 species).

The one family that is core in mouth and in gut but has a relatively narrow phylogenetic distribution (identified here in 33 species) is family #125536. This family codes for a 46-amino acid transmembrane protein. We hypothesize that this family is subject to horizontal transfer based on two observations (see section about horizontal transfer below). First, the small gene is found in the vicinity of genes that are known to participate in mobilization of DNA. Second, the family displays a sporadic distribution across multiple *Firmicutes* classes (Supplementary Table S5). Three of the families that are core to skin and were classified to multiple *Staphylococcus* species are homologs to beta class phenol-soluble modulin proteins (Supplementary Figure 6), a family of toxins that have multiple roles in staphylococcal pathogenesis (Cheung et al., 2014).

Altogether, our data suggests that among the small protein families there are families that may be important for the adaptation of the bacteria to a specific body niche and are probably not essential in other body niches.

### Small proteins that are potential mediators of cell-cell and cell-host communication

We were particularly interested in small proteins that could be involved in the crosstalk between microbial cells and their environment (host or other microbial cells). Communication is typically mediated through direct cell-cell contact or via small diffusible molecules secreted by cells (Hayes et al., 2010; Moreno-Gámez et al., 2017). We thus postulated that proteins that are at the cell surface or are secreted are more likely to be involved in cell-cell communication.

We looked in our dataset for small protein families that are either transmembrane and/or potentially secreted. To predict transmembrane and signal peptides, we applied two algorithms, TMHMM (Krogh et al., 2001) and SignalP (Petersen et al., 2011), on all 467,538 small proteins that constitute the 4,539 small protein families. We classified a family as transmembrane/secreted if ≥ 80% of the homologs of the family are predicted to be such. Due to the limitations associated with prediction of secreted proteins, we believe that the number of secreted proteins in our dataset is in fact higher than we predict here.

Of the 4,539 small protein families, 1,295 (28%) are predicted to be transmembrane, 35 (0.7%) small protein families are predicted to be secreted, and 29 (0.6%) families are predicted to be both transmembrane and secreted (Supplementary Table S5). Our prediction of secreted proteins relies on the presence of signal peptides, defined mostly from observations in the model organism *E. coli (Proteobacteria).* This bias is reflected in our analysis, in which 19/35 secreted small proteins are unique to the phylum *Proteobacteria*, that is known to actually be less abundant than other phyla (e.g. *Firmicutes* and *Bacteroidetes)* within the human microbiome.

To pinpoint small proteins that could be specifically important to life within the mammalian gut, we asked which of the transmembrane/secreted families have homologs in other mammalian guts but not in other niches (no other human body sites nor other non-mammalian metagenomes). Our mammalian gut metagenomes include 86 samples originating from diverse mammals, including mouse, rat, multiple non-human primates, panda and more (full list in Supplementary Table S6). This narrowed our set from 1,266 to 126 families (transmembrane = 122, secreted = 2, transmembrane and secreted = 2; Supplementary Table S5) that are found in human as well as other mammalian gut metagenomes.

Family #350024 drew our attention since it has the highest number of homologs in other non-human mammalian guts. We identified 30 homologs of this small protein in 13 different mammalian gut metagenomic samples. This family encodes for a 33-amino acid transmembrane protein with no annotated domain or known function. A homology search of the representative sequence of family #350024 against all 1,266 transmembrane families of the ~4k small protein families reveals that this small protein is actually even more abundant: there are 22 additional small protein families, ranging in size between 24-40 amino acids (Supplementary Table S8), that share sequence homology with this family, though they are divergent enough not to be clustered into one big protein family, suggesting rapid evolution (Figure 5A). These transmembrane proteins are often found in mammalian/bird gut samples and are in most cases encoded by diverse *Bacteroidetes* and *Firmicutes* species (Figure 5B).

**Figure 5.**
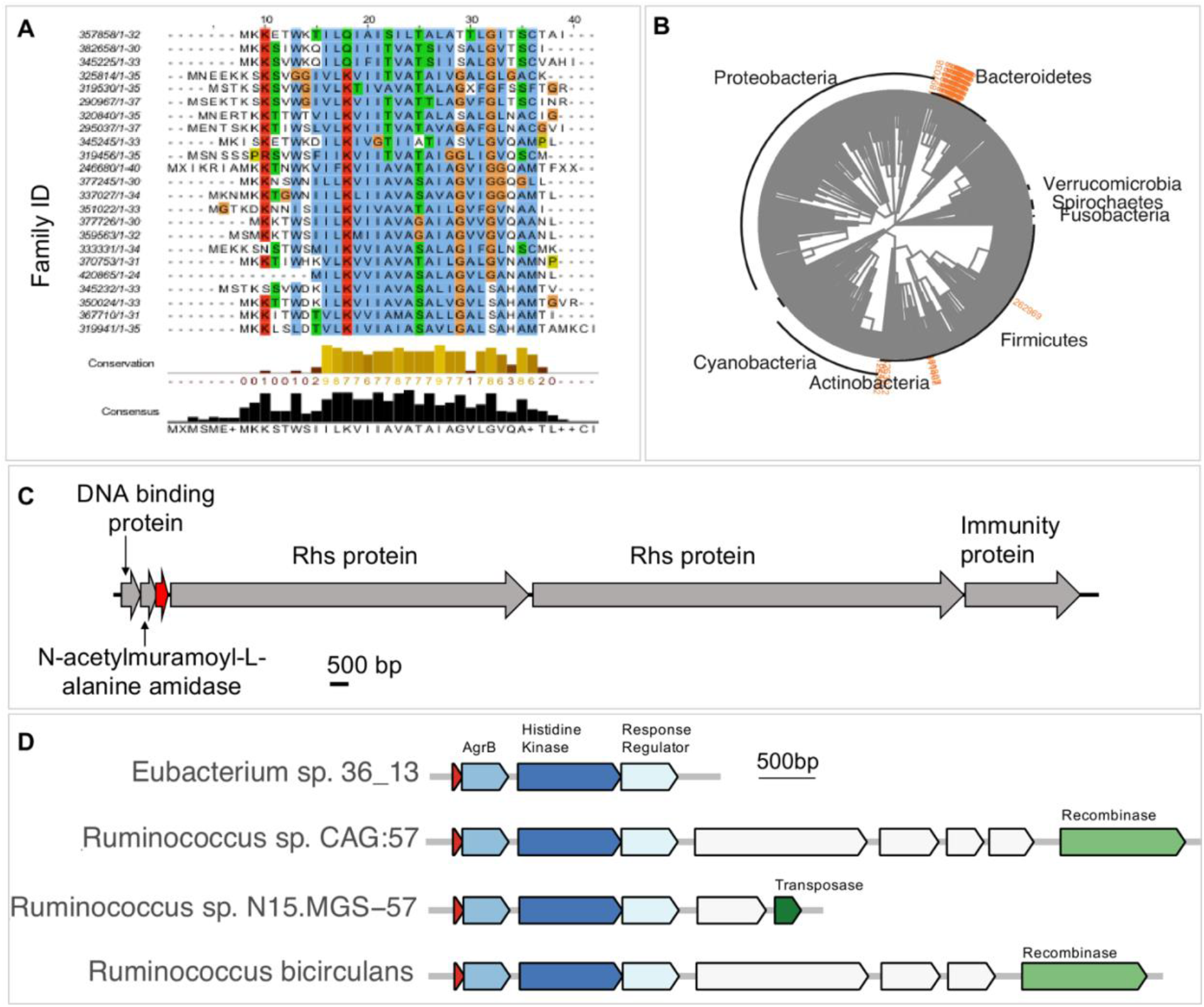
Small proteins that are potentially involved in cross-talk. **A-C**. Family 350024 is an abundant gut-related transmembrane family potentially involved in bacteria-host or bacteria-bacteria crosstalk. **A**. Multiple sequence alignment of representatives of all families that share amino acid sequence homology with family #350024. The length of the protein sequence is indicated after each family ID. **B**. Phylogenetic spread of family #350024 and 22 other homologous families. **C**. Genomic neighborhood, next to a DNA binding protein and an N-acetylmuramoyl-L-alanine amidase, an enzyme that cleaves the amide bond between N-acetylmuramoyl and L-amino acids in bacterial cell walls. The locus tag of the small transmembrane protein (red) is Ga0104402_10435 *(Bacteroides ovatus* NLAE-zl-C500). **D**. Putative novel signaling molecule that is presumably subject to horizontal transfer. Schematic representation of genes encoded on contigs of family #155173. In addition to Agr genes, these contigs typically harbor genes that are associated with horizontal transfer.

The genomic localization of this sORF is also conserved among homologs. Homologs of this small protein family are recurrently found adjacent to a DNA binding protein and an N-acetylmuramoyl-L-alanine amidase, an enzyme that cleaves the amide bond between N-acetylmuramoyl and L-amino acids in bacterial cell walls (Figure 5A). Interestingly, the product of an amidase was recently shown to mediate channel formation between bacterial cells that express them (Zheng et al., 2017). In addition, we often observe within the close vicinity to these three genes, virulence-related genes as VirE and/or genes encoding for the Rhs protein, a DNAse that is delivered to neighboring cells during contact dependent inhibition as well as the immunity protein that protects the encoding cell from the Rhs’s toxic effect (Koskiniemi et al., 2013) (Figure 5C). Altogether, we hypothesize that this small protein may be involved in crosstalk with other cells, potentially as part of a novel secretion/inhibition mechanism.

We were intrigued by the genomic neighborhood of family #155173, which was identified in over 40% of gut samples. Homologs of this 44-amino acid secreted protein are recurrently found upstream of three genes: a transmembrane protein annotated as AgrB, a histidine kinase and a response regulator (Figure 5D). This composition of genes strongly resembles the composition of the quorum sensing Agr operon, which consists of the short signaling peptide (AgrD), a transmembrane protein (AgrB), and a two component system composed of a histidine kinase (AgrC) and a response regulator (AgrA) (Olson et al., 2014). The small protein identified here was not assigned a domain in our query against CDD domains. However, the genomic localization of this secreted protein, next to AgrB and a two-component system, in addition to the similarity in size to AgrD, suggest that these four genes encode for a quorum sensing system, in which the signaling molecule component is a distant homolog/analog of AgrD. Intriguingly, we also observed in vicinity the presence of genes that mediate horizontal gene transfer, suggesting that this cluster of genes is subject to horizontal transfer (Figure 5D). The potential of the Agr quorum sensing system to undergo horizontal transfer has been suggested before, when homologs of the Agr genes were identified on the genome of a *C. difficile* temperate phage (Hargreaves et al., 2014), and here we provide additional support to this model.

### Small protein families with a potential role in bacterial defense against phage

The most abundant biological entity within the gut microbiome are viruses, the majority of which are bacteriophages that prey on bacteria (Mirzaei and Maurice, 2017). As a result, bacteria have evolved a variety of defense systems that protect them from phage attack, including restriction-modification systems, CRISPR, toxin-antitoxin and more (Dy et al., 2014; Koonin et al., 2017; Stern and Sorek, 2011). Defense genes tend to cluster in distinct mobile genomic regions, denoted ‘defense islands’ (Koonin et al., 2017). This notion has been recently used to identify multiple novel defense systems based on their localization within ‘defense islands’ (Doron et al., 2018).

Here, we were interested in identifying small proteins that could be associated with defense against phage. Small defense-related proteins are easily be missed in bioinformatic studies, such as the recent systematic study that aimed at identifying CRISPR-Cas related genes, which applied an inclusion cutoff of 100 amino acids (Shmakov et al., 2018), or studies that rely on domain annotation of protein families (Doron et al., 2018).

To identify small protein families that could be related to bacterial defense against phage, we identified the longest contig associated with each of the ~4k families (since short contigs code for fewer genes and are thus less informative) and screened them for the presence of known defense genes within ≤10 genes from the small protein. To identify defense genes, we used a list that was recently compiled by Doron et al. that contains 427 different COGs/Pfams of known defense genes (Doron et al., 2018). Based on this screen, we were able to identify 185 small protein families that may be associated with defense against phage (Supplementary Table S5). This group includes 27 small protein families that are encoded within the vicinity of CRISPR genes (Supplementary Table S5).

We took a more detailed look at two of the potential defense-related small proteins identified in our screen. Family #395508 is an example of a potential CRISPR-related small protein. It encodes for a 28-amino acid transmembrane protein identified here in multiple *Veillonella (Firmicutes)* species (Figure 6). In 65/72 (90%) contigs on which we detected homologs of this family, the small protein is encoded within ≤10 genes away from CRISPR-related genes. Another example of a potential defense related family is family #588. In 102/119 contigs within which we identified this small gene, the gene is encoded immediately upstream of a known ‘orphan’ toxin (i.e. devoid of a cognate antitoxin) that encodes a PIN nuclease (COG3744) (Supplementary Figure 7). Based on the ‘guilt by association approach’ (Leplae et al., 2011), we hypothesize family #588 codes for a novel antitoxin protein of a toxin-antitoxin system (i.e. not paired with ‘original’ antitoxins or toxins). Altogether, we provide a list of small proteins that are encoded within the vicinity of known anti-phage genes and as such represent an opportunity to identify novel defense genes that have, to date, escaped detection.

**Figure 6.**
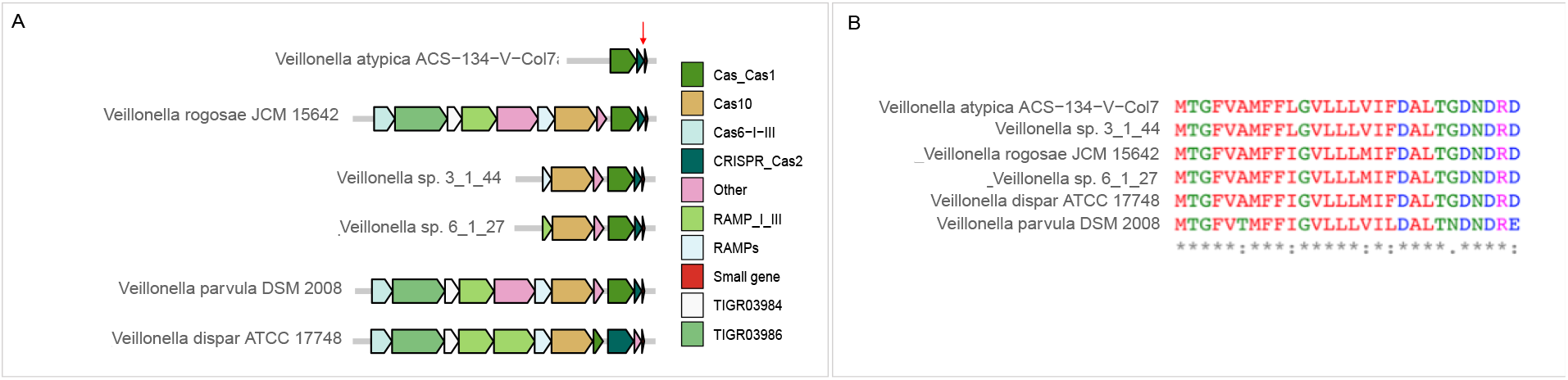
A small protein family (#395508) possibly asoociated with a CRISPR anti-phage system. **A**. Genomic neighborhood of small protein (red arrow) across 6 different species. Homologs of this small protein are shown in the genomic locus in which they were found among a variety of *Veillonella* species within HMPI-II data. **B**. Multiple sequence alignment of homologs of the family demonstrates a high level of conservation within small protein family #395508.

### Small proteins that are part of the ‘mobilome’

It has been suggested that the human gut serves as a “melting pot” of horizontal genetic material exchange, by which bacteria evolve to adapt to the ever changing conditions in the gut (Liu et al., 2012; Shterzer and Mizrahi, 2015). This phenomenon mediates transfer of antibiotic resistance genes, virulence genes, genes involved in metabolism and stress response, as well as genes involved in defense against phages (Ochman et al., 2000; Soucy et al., 2015; Zaneveld et al., 2008).

Here, we attempted to identify small protein families that could be part of the bacterial ‘mobilome’. A hallmark of genomic regions that are subject to HGT is the presence of genes that mediate horizontal transfer, including integrase and transposase (Oliveira et al., 2017). In addition, since horizontal transfer spreads genes between potentially distant bacterial lineages, genes that are subject to horizontal transfer may display a distribution that is discordant with the organismal tree of life (i.e. ‘patchy distribution’) (Cordero and Hogeweg, 2009). We used these two characteristics to identify a subset of genes that is potentially subject to HGT.

First, we used the longest contig in each family to search among the ~4k small protein families for those that are encoded in the vicinity (≤10 genes away) of genes that are known to mediate horizontal transfer (transposase/integrase, see ‘Methods’ section). This resulted in a set of 634 (14%, 634/4,539) small protein families (Supplementary Table S5). We characterized the phylogenetic distribution of these 634 families. Families that display a patchy distribution are more likely to be horizontally transferred. A patchy distribution is associated with families that are identified in a relatively small number of species across multiple clades (Figure 7A and Supplementary Figure 8).

**Figure 7.**
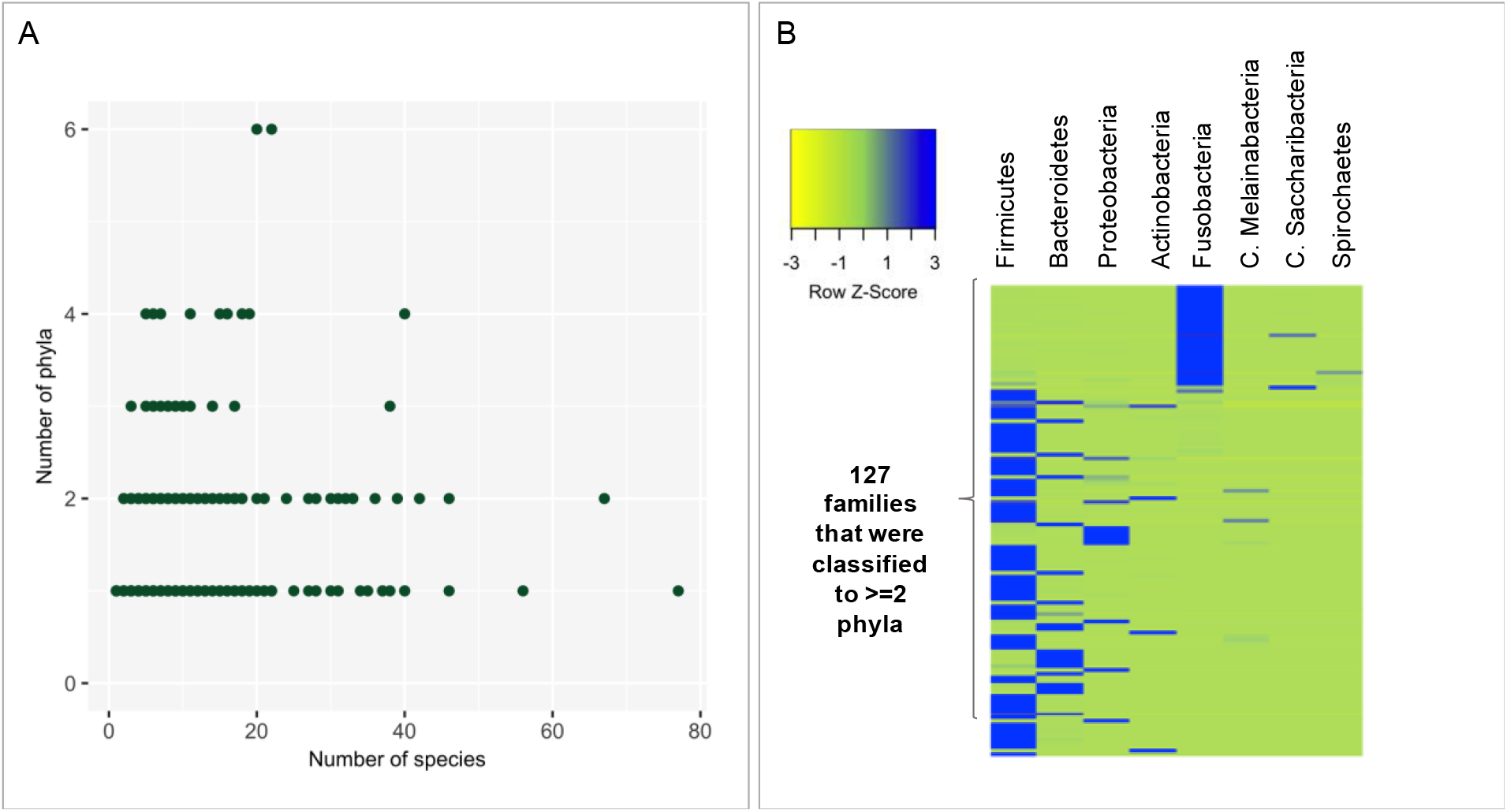
Small proteins that are potentially subject to HGT between phyla. **A**. Each dot represents one of 401 families that were identified in the screen of HGT genes in vicinity to small gene and whose median percentage of k-mers that were classified >=10%. Families that are encoded by a small number of species across a larger number of phyla are more likely to be true positives. **B**. Of the 401 families presented in A, 127 small protein families that were identified in ≥2 are presented. Only phyla that were identified in at least five different small gene families are shown. Each row was normalized.

It is generally assumed that HGT occurs more frequently between closely related organisms, but inter-phylum as well as inter-kingdom HGT events have also been documented, especially among organisms that share the same ecological niche (Caro-Quintero and Konstantinidis, 2015; Husnik and McCutcheon, 2018). Among the 634 small protein families identified in the initial screen as potentially horizontally transferred, we found examples of potential transfer within genus (e.g. family #380637 that was identified in 20 different *Staphylococcus* species), within family (e.g. family #12290 that was identified in 4 different *Ruminococcaceae* genera), within order (e.g. family #409298 that was identified in 8 different *Clostridiales* families), within class (e.g. family #420 that was identified in 3 *Actinobacteria* different orders), within phylum (e.g. family #222777 that was identified in 3 different Firmicutes classes) and even between phyla or between organisms belonging to different life domains. However, since a patchy distribution could be a result of sampling biases, our approach is more powered to detect HGT events between higher clades, such as between phyla. For a vertically transmitted gene to have a sporadic distribution across phyla, multiple deletion events of the gene across the tree should have occurred, which is less likely.

The 634 families that were identified in the screen for HGT-associated genes and were also mapped to multiple life domains and/or multiple phyla represent potential inter-phyla HGT events (Figure 7B). To enrich for small protein families in which the taxonomic classification is more reliable, we filtered out small protein families in which the median percentage of k-mers that could be classified is <10%, resulting in 401 small protein families. *Firmicutes* is the most represented phylum in this set, identified in 74% (297/401) of the families, in line with previous observations that this phylum is a major participant in genetic exchange (Caro-Quintero and Konstantinidis, 2015). Among the 401 small protein families there are 19 protein families that were also classified to viruses (Supplementary Table S5). These may represent potential transduction events, in which bacteriophages mediate the horizontal transfer of DNA between hosts.

## Discussion

Accumulating evidence suggests that small proteins play important roles in bacterial physiology. However, due to computational and experimental limitations, this class of proteins is consistently overlooked. Here, we focused on small proteins encoded by the human microbiome. We were interested in small proteins within this niche for several reasons. In terms of size, small proteins can represent a ‘bridge’ between the small molecule (natural product) world, a rich source of biologically active molecules such as antibiotics, and the larger protein world. As such, they are likely to display a range of activities that would resemble either class and thus operate at microbe-host interface. While natural products have attracted much attention and investigation (Donia et al., 2014; Milshteyn et al., 2018; Trivella and Felicio, 2018; Wilson et al., 2017), and large proteins are easier to detect and analyze, small proteins in the human microbiome have thus far evaded thorough systematic analysis. Identification of small proteins within the human niche would advance our understanding of this complex ecosystem and could present an untapped source of novel therapies. Moreover, it has the potential to reveal new biological insights about proteins of this size.

In this study we applied a combination of computational approaches on 1,773 healthy human metagenomes and identified a set of nearly 4,539 small protein families, that are with high confidence, not spurious. We show that most are unknown, that is not represented in traditional reference genomes and/or do not contain a known protein domain. By classifying the protein families according to their taxonomic distribution, their prevalence across human body sites and non-human metagenomes, their predicted cellular localization, and their genomic neighborhood, we assign putative functions to a subset of the families. To the best of our knowledge, this is the first exhaustive characterization of small proteins encoded by the human microbiome. This study only scratched the surface of what seems to be the full potential hidden in the human-associated small microbial protein space. We expect that data presented and the approaches used in this study will be an important resource and motivation for future studies.

Proteins that play housekeeping roles are expected to be unrelated to a specific niche or taxonomic clade. Here, through taxonomic classification of encoding contigs and classification of contigs according to their niche of origin (human body sites as well as diverse ecological niches) we identify eight small protein families that we predict to be housekeeping proteins. The exact functions of these highly conserved protein families and their potential relevance to the human host is yet to be discovered.

We focus on one of the predicted housekeeping families, that contains a domain of unknown function 4295 (DUF4295), and show that it is likely to be a novel small ribosomally associated protein, which may interact with the ribosome directly or indirectly. One may wonder how such a protein might escape detection, as ribosomes have been subject of deep investigation spanning several decades of research. We believe that this is due to the focus of prior research on a handful of model organisms and the dismissal of small ORFs from bioinformatics analysis pipelines. Bacterial ribosomes have mostly been studied in model organisms, such as *E. coli*, which lacks this predicted small protein. Many of the genomes that encode this small protein are residents of the human microbiome, whose genomes have mainly been sequenced in the last decade and whose ribosomes have not been studied, in depth. The experimental laboratory strain *Bacteroides thetaiotaomicron* VPI-5482 encodes this small proteins but due to its small size, the gene that encodes for this protein remained unannotated in the *Bacteroides thetaiotaomicron* VPI-5482 reference genome. Herein, our metagenomic analysis approach corrects for this bias to reveal a plethora of previously uncharted small protein families including this putative novel ribosomally associated protein.

Because of their short length, small proteins generally consist of one domain and represent a useful model system for protein folding simulations (Imperiali and Ottesen, 1999) and drug design (Vita, 2000). A detailed protein analysis study has recently suggested that the number of domains is reaching saturation (Scaiewicz and Levitt, 2018). However, more than 95% of the small protein families identified in our study do not significantly resemble any previously known domains. This stresses the possibility that the space of small proteins represents an untapped opportunity for discovery of new building blocks of proteins. Many studies rely on domains assigned to proteins. For example, two recent papers that systematically searched for genes associated with phage defense systems based their search on domains of known defense systems (Doron et al., 2018; Shmakov et al., 2018). This emphasizes the need to annotate domains of small proteins, or they will continue to be missed in future studies of this type, thus limiting our knowledge about entire classes of proteins such as phage defense systems, for example.

Transmembrane and secreted proteins mediate most of the interactions of a bacterium with its environment, making these classes of proteins relevant targets for medical research and potential drug targets. Here, we identified 126 transmembrane/secreted small protein families that were identified exclusively in mammalian guts (human as well as other mammalians). One of the families that is presumably very abundant across different mammalian guts encodes for a transmembrane protein that is encoded between a DNA binding protein and an amidase enzyme that cleaves cell wall. A recent paper showed that a similar enzyme is involved in formation of channels for material exchange between cells (Zheng et al., 2017). We suggest that the small protein identified is part of a cluster of genes that could also be involved in channel formation between cells and subsequent DNA translocation.

While HGT events within bacteria and archaea are unequivocal (Soucy et al., 2015; Wagner et al., 2017), the frequency and importance of HGT between domains of life remains less clear (Husnik and McCutcheon, 2018). Using taxonomic contig classification, we identified multiple families that were mapped to more than one domain of life, including 122 families that were mapped to both bacteria and eukaryotes and 30 families whose underlying contigs were mapped to both bacteria and archaea. While misassembly or misclassification of contigs could possibly account for this, this observation remains intriguing as it suggests either ancient conservation of sORFs or true genetic transfer between evolutionarily distant organisms.

Despite the power of the approach described in this work, we note a number of limitations. First, our analysis filters out families if they are encoded by < 8 different sequences. We are therefore liable to miss genuine small protein families when these are very rare. In addition, small proteins undergoing rapid evolution may fall into separate families in the sequence based clustering step, which may lead to them being filtered out due to small family size. Since we conduct our analysis on contigs and not on reference genomes, we are also vulnerable to errors in taxonomic classification that could stem from misassembly and/or misclassification of contigs. Next, our prediction of secreted proteins relies on the presence of signal peptides. This is a limited prediction from two opposing directions: first, not all proteins that harbor a signal peptide are secreted outside of the cell, as is the case in Gram-negative bacteria that require an additional mechanism to secrete the proteins outside the cell. Second, proteins that do not contain a signal peptide could still be secreted by other secretion machineries (Green and Mecsas, 2016), for which the sequence features are not well defined. In addition, signal peptides contain a hydrophobic region that can be mistaken for a transmembrane region, implying that a subset of the predictable transmembrane proteins could actually be secreted (Krogh et al., 2001). Finally, our analysis of the genomic region of small genes is limited by the number of genes that are encoded on the encoding contig, which is in turn limited by the length of the contig (which is, by itself variable in metagenomic data).

To advance from this study, mechanistic studies will be required. Gene deletion and complementation studies are likely to be highly informative, when studying small proteins genetically tractable bacteria, such as *Bacteroides thetaiotaomicron.* Alternatively, in light of their small size and the relatively low cost of their synthesis, it may be feasible to conduct high-throughput studies in which a selected subset of small genes is synthesized and heterologously expressed within cells to study gain of function phenotypes. Small proteins that are predicted to be secreted could be synthesized directly and applied on cells (bacteria or host) externally to identify novel mechanisms of cell-cell communication. As proteomic studies of human microbiomes are emerging, it is important to have comprehensive protein databases, since proteins that are not annotated in databases cannot be identified in mass spectrometry-based proteomics analyses. Thus, our study highlights the importance of optimizing experimental workflows for protein analysis and annotation to capture small proteins.

Knowledge of small peptides encoded by human gut bacteria is very limited. We hope that the data and computational approach presented here will open a new frontier in the study of the microbiome and its implications in health, while also providing a rich resource for both the basic science and translational medicine research communities.

## Materials and Methods

### Identification of sORFs from multiple human associated metagenomes

Contigs from 1,773 HMPI-II human-associated metagenomes from 17 body sites that were shotgun sequenced and contained no less than 5M bp sequenced per sample were downloaded from https://www.hmpdacc.org/hmasm2/. Body sites were collapsed into four groups (Supplementary Table 2). For each metagenomic sample, all ORFs were predicted using MetaProdigal (Hyatt et al., 2012) with parameters adjusted to include ORFs ≥15bp. Small ORFs where filtered to include only those that contain a start and stop codon, resulting in a set of 2,514,099 sORF ≤150bp.

### Clustering of sORFs into families

Proteins encoded by this set of sORFs were clustered using CD-Hit with the following parameters: -n 2 -p 1 -c 0.5 -d 200 -M 50000 -l 5 -s 0.95 –aL 0.95 –g1 (the shorter sequences were required to be ≥95% length of the representative of the cluster and the alignment must cover ≥95% of the longer sequence). This resulted in 444,054 clusters. Each cluster was assigned a ‘cluster representative’ by CD-Hit that was used in subsequent parts of our analysis.

### Domain Analysis

The CDD DB was downloaded from ftp://ftp.ncbi.nih.gov/pub/mmdb/cdd/little_endian/Cdd_LE.tar.gz on October 2018. This DB contains models that are in the default CDD database: CD (alignment models curated at NCBI as part of the CDD project), Pfam, Smart, COG, PRK and TIGRFAM. The amino acid sequence of each of the cluster representatives from each one of the 444,054 clusters was searched against this DB, using RPS-blast. A hit was considered significant if the e-value ≤ 0.01 (default CDD e-value threshold) and the small protein aligns to at least 80% of the PSSM’s length. Small protein families were classified according to the PSSM they hit. The same domain may be assigned to multiple families reflecting distant sequence conservation across families (e.g. families #263535, 73615 and 209227 that were all assigned the same PSMM of a small ribosomal protein L36), and families may be assigned multiple domains, reflecting redundancies in CDD (e.g. family #305829, that was assigned two different PSSMs, both of ribosomal protein L34).

### Identification of known proteins among the small protein clusters

Small proteins that were studied in depth were divided into two groups: those that have an assigned sequence domain and those that do not. The domains that were assigned to the 444,054 clusters were queried for domains of the first group. The known small proteins that do not have an assigned sequence domain were queried against all 444,054 representative protein sequence, using blastp with word-size 2. Hits were considered significant if: e-value ≤ 0.05, the alignment spans ≥90% of the protein and the length of the hit was 90%-110% of the length of the small protein.

### Assigning p-values to small protein families

RNAcode (Washietl et al., 2011) was applied on all 11,715 clusters that are composed of ≥8 different DNA sequences. Only the 4,543 that were assigned a *p*-value of ≤ 0.05 were analyzed in subsequent steps of analysis.

### Taxonomic classification of small protein families

Classification of 1,504,527 contigs encoding the small proteins were classified using the One Codex database 2018. Each contig was compared to a database of 83,701 microbial genomes. microbial reference genomes. The platform matches all overlapping k-mers in a given contig to the most specific organism possible. Since not all k-mers are unique to a specific operational taxonomic unit (OTU), each k-mer was classified to the lowest common ancestor (LCA). Individual k-mer matches across a given contig were then aggregated to assign the most specific and consistent OTU to the contig. For each contig, the proportion of 31-mers that were classified out of the total 31-mers (rounded to the nearest whole number), was recorded. For each small protein family, the number of different OTUs, phyla, classes, orders, families, genera and species in which it was detected was recorded. Of the 1,504,527 total contigs, 69,974 contigs (4.6%) could not be taxonomically classified. The four families in which all contigs were classified as *‘Homo sapiens*’ were excluded from further analysis.

### Analysis of small proteins in RefSeq genomes

Protein sequences from 69,681 RefSeq genomes, were download from ftp://ftp.ncbi.nlm.nih.gov/genomes/RefSeq/bacteria/ on July 2017. Representative protein sequences of 4,539 families were blasted against 3,549,250 RefSeq proteins that are ≤ 50 amino acids with word_size 2 and Max number of hits = 500. To call for small genes on these RefSeq genomes, Prodigal (Hyatt et al., 2010) with parameters adjusted to include ORFs ≥15bp was run on RefSeq reference genomes and representatives of families were blasted against 6,931,965 prodigal-predicted proteins that are ≤50 amino acids as described above. In both cases, hits were considered significant if: e-value ≤0.05, the alignment spans ≥90% of the query protein and the length of the hit is 90%-110% of the length of the small protein.

### Analysis of genomic neighborhood of small proteins

MetaProdigal (Hyatt et al., 2012) was used to call for genes on longest contig of each of the families. Amino acid sequence of all the genes on all longest contigs were searched against CDD, using RPS-blast with an e-value threshold of 0.01. A hit was considered significant if the e-value ≤ 0.01 (default CDD e-value threshold) and the protein aligns to at least 80% of the PSSM’s length. Each gene on a contig could have multiple significant domains. All domains of genes identified within 10 genes away from the small gene were recorded. For example, for family #111917, the small gene on the longest contig associated with the families is the 14^th^ gene on the contig. The list of genes in vicinity includes “10: Phage_holin_4_1, COG4824, holin_tox_secr”, meaning that the 10^th^ gene on the contig was assigned these CDD domains.

To identify small proteins that are in the vicinity of defense genes, domains identified within ≤10 genes away of the small protein were queried against the list of COGs and Pfams downloaded from Supplementary Table S1 in Doron et al. (Doron et al., 2018) as well as against the words ‘CRISPR’, cas1 and cas2. For families that were found positive in this step, all contigs of the families were annotated and queried against CDD as described for the representative contig. Contigs in which defense genes were identified within 10 genes away from the small gene homolog were classified using One Codex (described above) and recorded (Supplementary Table S5). To identify HGT-related contigs, the words ‘recombinase’, ‘integrase’, ‘transposon’ and ‘transposase’ were queried. To identify contigs that are presumably prophage, the words ‘phage’, ‘terminase’, ‘tail’, ‘caspid’ and ‘portal’ were queried against the CDD ‘short’ domain description. Only families in which the word ‘phage’ as well as one other others in the list were identified, were recorded as ‘phage’. Supplementary Table S5 lists all domains found on longest contigs of each family.

### Identification of homologs of family #350024

Protein sequence of representative of family #350024 was blasted against all 1,295 amino acid sequences of representatives of transmembrane families, using a *p*-value threshold of 0.05. For each of the homologous families, all contigs associated with the family were annotated using MetaProdigal and proteins on all contigs were subject to RPS-blast against CDD database.

### Identification of species that encode for the small protein adjacent to known toxin (families #855 and 223481)

All contigs of family #855/#350024 were annotated with MetaProdigal, proteins were queried against CDD to assign domains. Contigs in which the small gene is encoded immediately upstream/downstream of PIN domain/COG3943 were classified using the One Codex database, as described above. Number of species in which the small gene and the PIN domain were identified one next to the other, was recorded.

### Mapping of small proteins to body parts

For each member in each of the ~4k families, we recorded the patient and body site from which it originated. A member of a small protein family that was detected more than once in a specific body site of a specific patient was counted only once (even when identified in multiple visits). For every family, the total number of appearances in each type of body site was then calculated. The total number of body samples from a specific body site counts multiple samples from the same patient’s body site, as one.

### Search against non-human metagenomes

DNA sequences of each of the cluster representatives was blasted against a set of 5,829 non-human metagenomes using blastn with e-value 1e-05, 50% identity and alignment length coverage of 90%. MetaProdigal, adjusted to small gene finding (see above) was applied on the contigs that were hit in the previous step. The proteins that were identified by prodigal were then used as a DB against which protein sequences of all representatives was blasted against. Hits were considered significant if: e-value ≤ 0.05, the alignment spans ≥90% of the query protein and the length of the hit is 90%-110% of the length of the small protein.

### Cellular Localization

SignalP was run with default parameters once with ‘gram +’ and once with ‘gram -’ mode on all small proteins encoded by ~4k families. TMHMM was run on the same set of proteins with default parameters. For every family, and for every attribute (transmembrane/signal ‘gram +’/signal ‘gram -’) the number of transmembrane helices was counted, and whether the protein is predicted to be secreted. The percentage of family members that were predicted to be transmembrane/secreted was calculated and a family was considered transmembrane/secreted if ≥80% of the family members were predicted to be such.

## Supporting information

## Acknowledgments

We thank all members of the Bhatt laboratory for providing feedback on the study design, bioinformatics pipeline and manuscript revisions. Special thanks to Brayon Fremin and Soumaya Zlitni from the Bhatt lab for fruitful scientific discussions. We thank Noam Livnat, Asaf Levy and Oren Kolodny for useful discussions and insightful comments throughout the course of this study. We thank Amiyaal Ilany for reading and commenting on the manuscript. We also thank Ramesh Nair for assistance in downloading of data. We thank Andrea Scaiewicz for insightful discussion about protein domains. This work was supported in part by PhRMA Foundation (H.S.) and by the NIH/HNGRI T32 HG000044 grant (H.S.). A.S.B. was funded in part by the National Cancer Institute NIH K08 award, no. CA184420 and the Damon Runyon Clinical Investigator Award. The content is solely the responsibility of the authors and does not necessarily represent the official views of the NIH.

## Competing interests

The authors declare no competing interests.

## Corresponding author

Correspondence to Ami S. Bhatt

