## Supplementary material for "Large-scale analyses of human microbiomes reveal thousands of small, novel genes and their predicted functions"

*Supplementary Notes:*

This file contains additional information about known proteins in HMPI-II data, about small proteins of phage origin, about small protein families that are identified across multiple body sites, about non-bacterial small proteins and guidelines for extraction of contigs associated with specific family and supplementary figures.

*Supplementary Tables:*

**Supplementary Table S1** List of 1,773 HMPI-II samples used in this study.
**Supplementary Table S2** Number of samples from each body site.
**Supplementary Table S3** List of 444,054 clusters
**Supplementary Table S4** Representation of 29 known small proteins in our database. The first tab lists the hits to known small proteins that have an assigned domain. The second tab lists the hits to known small proteins that do not have an assigned domain or failed the domain search and where queried using BLASTp.
**Supplementary Table S5** List of 4,539 small protein families
**Supplementary Table S6** List of 5,829 non-human metagenomes used in this study
**Supplementary Table S7** Number of contigs classified to different taxonomic clades by body site
**Supplementary Table S8** List of families that are homologous to family #350024

*Supplementary Files:*

**Supplementary File 1** Amino acid sequences of all ~4k families.
**Supplementary File 2** Nucleotide sequences of all ~4k families.
