## Supplementary material for "Large-scale analyses of human microbiomes reveal thousands of small, novel genes and their predicted functions"

*Only a small subset of well characterized small proteins are relevant to the human microbiome*

Another example of a small protein that was extensively studied and also happens to be present in the human microbiome is ComC, which is a quorum-sensing signal that enables *Streptococci* to regulate DNA uptake and genetic transformation in response to population density as well as environmental queues such as antibiotic stress (Moreno-Gámez et al., 2017). We found ~2% (2,176/113,693) small proteins, clustering into 19 clusters were homologous to ComC. Once again, the relatively large number of clusters may reflect the genetic variability that has previously been associated with this protein (Allan et al., 2007).

Less abundant but still present are the CydX (YbgT) domain, a small protein required for the function of cytochrome *bd* oxidase (Sun et al., 2012); KdpF, part of the high-affinity ATP-driven potassium transport system (Gaßel et al., 1999), the toxins Hok (Chukwudi and Good, 2015) and TisB (Steinbrecher et al., 2012) as well as the multidrug efflux pump accessory protein, AcrZ (Hobbs et al., 2012) and SgrT, regulator of glucose metabolism (Lloyd et al., 2017).

For the known small proteins that do not have an assigned domain and for those that failed the domain search, we used BLASTp (Camacho et al., 2009) to search for their sequence against all representatives. We identified a small number of homologs of MntS, that takes part in manganese chaperoning (Martin et al., 2015); PmrR, regulator of a membrane-bound enzyme (Kato et al., 2012); SidA, inhibitor of cell division (Modell et al., 2011); MgtS (formerly known as YneM), modulates intracellular $Mg^{2+}$ levels to maintain cellular integrity upon $Mg^{2+}$ limitation (Wang et al., 2017), and Blr, involved in B-lactamase resistance (Karimova et al., 2012).

We were, however, not able to identify small proteins with the MciZ (Handler et al., 2008), MgrB (Salazar et al., 2016), SpoVM (Cutting et al., 1997) or SdA (Rowland et al., 2004), FbpA/B/C (Gaballa et al., 2008), MgtR (Choi et al., 2012), Pril42 (Impens et al., 2017), PepA1 (Sayed et al., 2012), BacSp222 (Wladyka et al., 2015), Listeryolysin S (Quereda et al., 2017), Streptolysin S (Molloy et al., 2011) and AimP (Erez et al., 2017) within our dataset of human associated metagenomes.

*Small proteins of phage origin*

A growing number of studies show that phages play important roles beyond preying on bacteria. In addition to being agents that mediate horizontal transfer of genes between hosts, viral genomes often contain genes that originate from host cells (auxiliary genes), that provide fitness advantage to these viruses upon their expression (Breitbart et al., 2003; Colomer-Lluch et al., 2011; Hurwitz et al., 2016; Manrique et al., 2017; Virgin, 2014). Since phage genomes are considered to be 'compact' (i.e. non-beneficial genes are lost through selective evolution) (Brüssow and Hendrix, 2002), discoveries of bacterial gene homologs within phage genomes hold promise for interesting discoveries.

We thus sought to identify small proteins that have homologs among phage genomes, and as such could represent novel auxiliary genes. We started by classifying all small-protein encoding contigs against a database that included a set of 19,879 viral genomes (Supplementary Table S8). We were able to identify 182 families in which at least one contig was classified as 'viral' (Fig 7A and Supplementary Table). By far the most common phylogenetic distribution that includes a viral component, observed in 161 (161/182, 88%) of the viral small protein families, is of bacteria-virus (Fig 7B and Supplementary Table).

Contigs that harbor prophages, bacteriophages integrated into the host's genome, could theoretically be classified as either 'bacterial' or viral, depending on whether the viral reference genome database contains their sequence and on the relative part of sequence that is of viral origin on the contig. To predict prophage regions that were classified as bacterial, we used a common, complementary approach that is based on detection of known viral genes, such as the terminase, capsid, tail and portal proteins (Roux et al., 2015). We screened for a list of 'phage genes' encoded on longest contigs of families (Methods). We were able to identify 223 of the 4,539 small protein families in which none of the homologs-encoding contigs was classified as 'viral', but nevertheless the longest contig encodes for at least one phage gene (Supplementary Table S5 and Methods). Altogether, we identified 405 small protein families that either have at least one homolog that was classified as viral or are integrated within a presumable prophage region (Supplementary Table S5) (147 families that were classified as viral, 223 families in which we identified phage genes and 35 that were both classified as viral and the longest contig also harbors signature phage genes).

*Small protein families that have a broad distribution across body sites*

In our search for small proteins that have a 'general' rather than niche specific function, we sought to identify families that are not associated with a specific body niche. To do so we mapped all contigs of all families back to one of the body sites (mouth, gut, skin or vagina) from which it was assembled (Figure 1A) and asked which families consist of contigs that were mapped to all of these body sites.

Among the ~4k families, we identified 55 (55/4,539, 1.2%) that have a 'broad' distribution across body sites. This is probably an underestimation since our samples are biased against skin and vagina samples (Supplementary Table S2), making it less probable to identify small proteins in these two niches. Seven of these families are families that are also very abundant taxonomically (i.e. among the 14 families that have homologs in ≥100 species).

For the remaining 48 families (55-7), we were able to assign a CDD domain to only 16% (8/48) of these families. Among these are six families that encode for ribosomal proteins, the rubredoxin domain (participates in electron transfer), and a family assigned a DUF3687 domain.

Altogether, using taxonomic classification and assignment of homologs to body site of origin, we were able to identify a handful of small protein families that could be playing key roles based on their widespread spatial and taxonomic distribution.

*The non-bacterial fraction of the small protein families*

Disregarding the unclassified homologs, the vast majority of protein families in this set are classified as bacteria (4,189/4,539, 92%) (Supplementary Table S5). In addition, there are 8 families that are classified to Eukaryotes and 152 families that are classified to multiple life domains (Supplementary Figure 9). For example, family #241192 was classified to two life domains (bacteria, eukaryote) as well as to virus. In the contig that was classified to Eukaryotes, ~50% of the *k*-mers were classified as *Candida albicans,* an opportunistic pathogenic fungus, common in the human microbiome (Sam et al., 2017) (Supplementary Table S5).

*Guidelines for extraction of all contigs associated with a specific family*

For the sake of this explanation, family #314163 was chosen. There are 8 homologs in this family.

1. In the Supplementary file that contains all family sequences, retrieve all sequences that follow the header "Family: 314163". This results sequences listed below.
2. For each sequence, retrieve the field that indicates the sample number and contig number (both in bold in the sequences below) according to the following convention:

>familyID*_*Taxon ID of underlying contig*_*Name of Taxon*_*percentage of 31-mers that were classified*_*Body site from which contig originated **_SampleID**_PatientID_Female/Male_Visit nummer_"prediction"_contig number_locationOfSmallGeneOnContig
3. Download the relevant sample, according to the Sample ID, from https://www.hmpdacc.org/hmasm2/.
4. Retrieve the contig of interest from the sample file, according to the contig number, which was given by HMPI-II.

In this example, for the first homolog of family #1179, download the sample SRS06477 and retrieve contig number 22054. This contig codes for the small gene. The small gene is the third gene, if genes are called on this contig with MetaProdigal with parameters modified to include all ORFs as short as 15bp.

Family: 314163
>314163*_*32207*_*Rothia*_*97%*_*Tongue_dorsum_SRS064774_764062976_Female_2_prediction_22054_3
MXTHKRLIDVVDPTPKAVDALMRLDLPADVNIEIKL
>314163*_*1120941*_*Actinomyces dentalis DSM 19115*_*81%*_*Supragingival_plaque_SRS074598_159733294_Female_2_prediction_5243_2
MRTHKRLIDIIDPTPKAVDSLMRLDLPADVNIEIKL
>314163*_*1120941*_*Actinomyces dentalis DSM 19115*_*82%*_*Supragingival_plaque_SRS077520_764285508_Male_2_prediction_contig-100_3567.148361_1
MRTHKRLIDIVDPTLKTIDTLRRLDLPADVNVEIKL
>314163*_*1739431*_*Rothia sp. HMSC061E04*_*92%*_*Tongue_dorsum_SRS1055059_316129862_Female_1_prediction_9420_11
MRTHKRLIDVVDPTPKAVDALMRLDLPADVNIEIKL
>314163*_*29465*_*Veillonella*_*98%*_*Supragingival_plaque_SRS143684_868454789_Male_2_prediction_9536_6
MRTHKRLIDILEPNSKTVDAITRLDLPAGVSIEIKL
>314163*_*32207*_*Rothia*_*82%*_*Supragingival_plaque_SRS147106_147406386_Male_3_prediction_47966_3
MRTHKRLIDVVDPTPKAVDALMRLDLPADVNIEIKL
>314163*_*32207*_*Rothia*_*95%*_*Supragingival_plaque_SRS149237_901775393_Male_2_prediction_contig-100_7304.58295_2

MRTHKRLIDVVDPTPKAVDALMRLDLPADVNIEIKL
>314163*_*1125718*_*Actinomyces massiliensis F0489*_*86%*_*Supragingival_plaque_SRS893280_486505039_Male_1_prediction_315_2
MRTHKRLIDIVDPTPKAVDSLMRLDLPADVNIEIKL

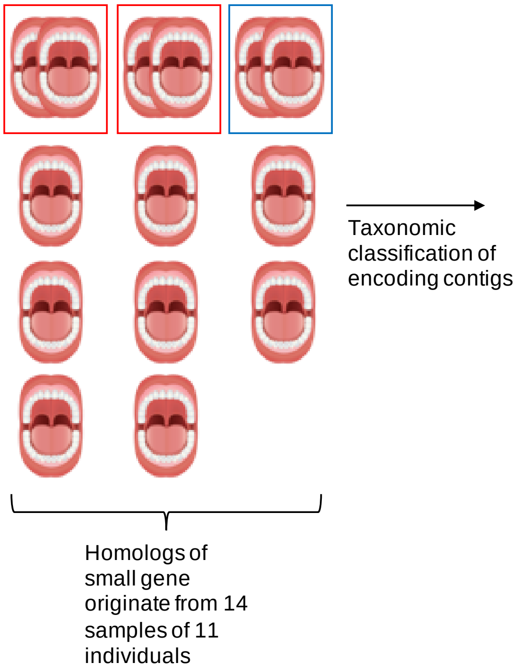


| Origin of Contig | Taxonomy | Individual | Transmembrane? |
|---|---|---|---|
| Sample 1 | Unclassified | 1 | Y |
| Sample 2 | Unclassified | 1 | Y |
| Sample 3 | Unclassified | 2 | Y |
| Sample 4 | Unclassified | 2 | Y |
| Sample 5 | *Actinomyces naeslundii* | 3 | Y |
| Sample 6 | *Pepy6virus* | 4 | Y |
| Sample 7 | Unclassified | 5 | Y |
| Sample 8 | Unclassified | 6 | Y |
| Sample 9 | Unclassified | 7 | Y |
| Sample 10 | Unclassified | 8 | Y |
| Sample 11 | Unclassified | 9 | Y |
| Sample 12 | Unclassified | 10 | Y |
| Sample 13 | Unclassified | 11 | Y |
| Sample 14 | Unclassified | 11 | Y |

**Supplementary figure 1. Explanation of numbers associated with each family as they appear in supplementary tables, through family #221403.** Small proteins of this family were identified 14 times across samples (hence, number of members in cluster = 14) originating from 14 metagenomic mouth samples that were sampled from 11 individuals: in 2 individuals the small gene was identified in two subsequent visits (red box) and in 1 individual the small gene was identified in two samples of two different mouth sublocations (Supragingival_plaque and Subgingival_plaque), that were taken from the same individual at the same visit (blue box). To avoid redundancy, we counted each individual only once, hence ‘Number of times found in Mouth’ = 11. Classification of each of the 14 contigs resulted in 3 OTUs, as all ‘Unclassified’ were counted as one single OTU, in lack of additional information, hence ‘Number of OTUs’ = 3. ‘Number of Bacterial species = 1’ since only one of this is a bacterial species and ‘Number of non-bacterial OTUs’ = 1 (in this case Pepy6virus’). All small proteins in this family have a predicted transmembrane domain, hence ‘% of family members that are predicted to have a transmembrane domain’ = 1.

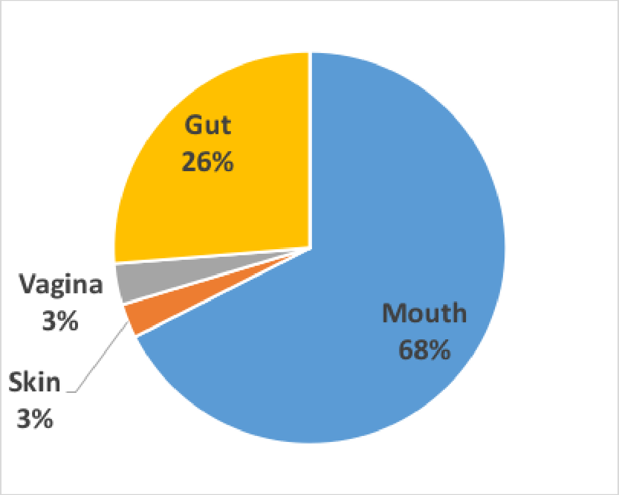


**Supplementary figure 2. Proportion of samples belonging to different body sites.** 128,368,337 contigs were obtained from 1,773 HMPI-II human-associated metagenomes that were shotgun sequenced, spanning 4 different major body sites from 263 healthy individuals. Samples were obtained from individuals in one, two or three subsequent visits. Body sites were collapsed into four groups: anterior nares, buccal mucosa, hard palate, keratinized gingiva, palatine tonsils, saliva, subgingival plaque, supragingival plaque, throat and tongue dorsum to 'mouth'; mid vagina, posterior fornix and vaginal introitus to 'vagina'; left retroauricular crease, right retroauricular crease and right antecubital fossa to 'skin'; stool samples were renamed 'gut' here.

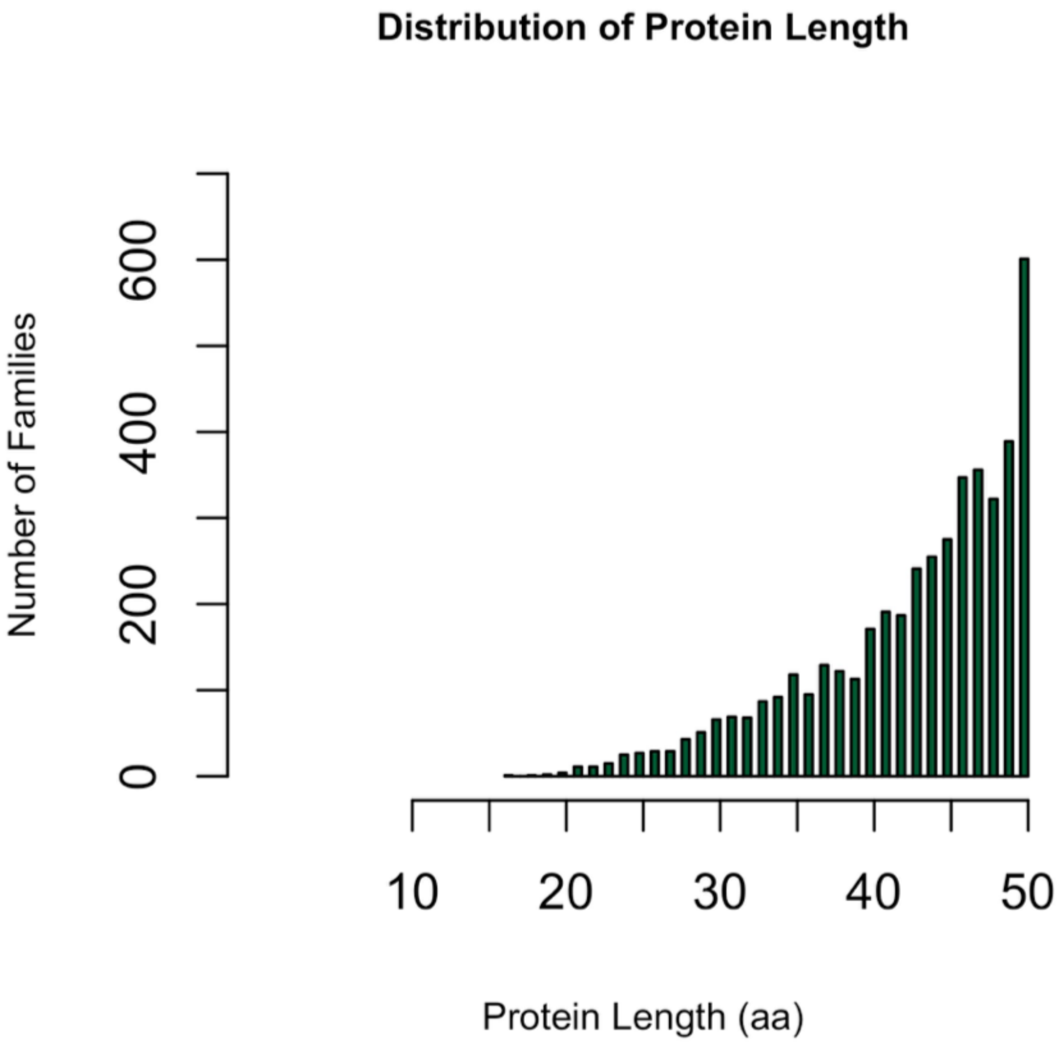


**Supplementary figure 3. Protein length distribution of 4,543 families.** Proteins of >50 aa or <5 aa were filtered out. The smallest protein in our dataset is encoded by family #442207 and is 16aa long.

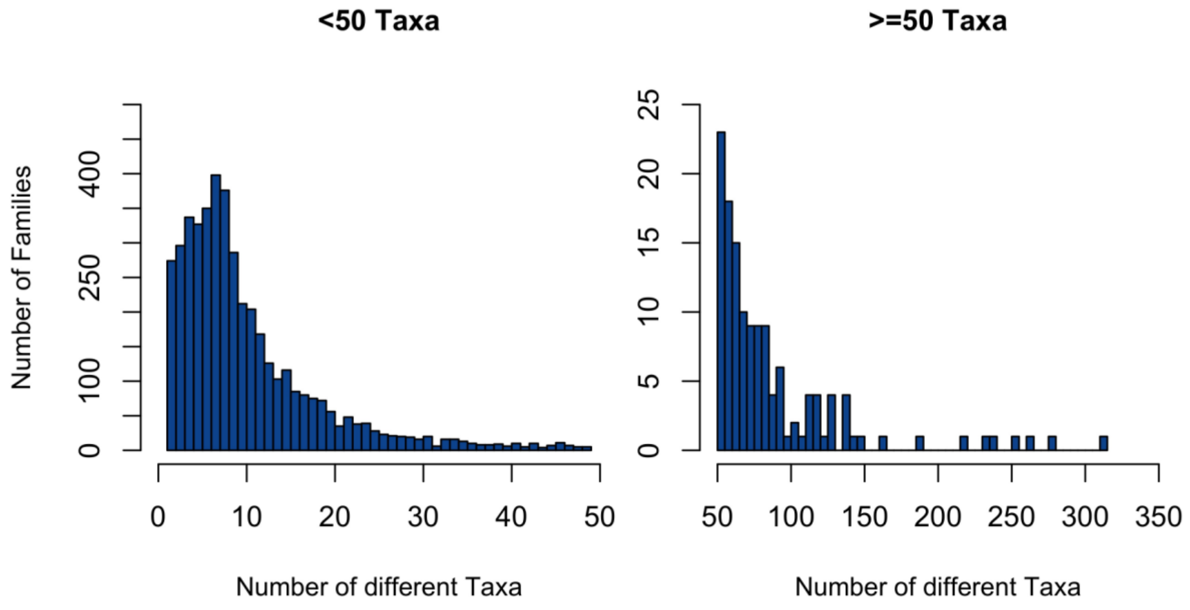


**Supplementary figure 4. Distribution of number of encoding taxa per family. If multiple homologs are mapped to the same taxa, they were counted only once.**

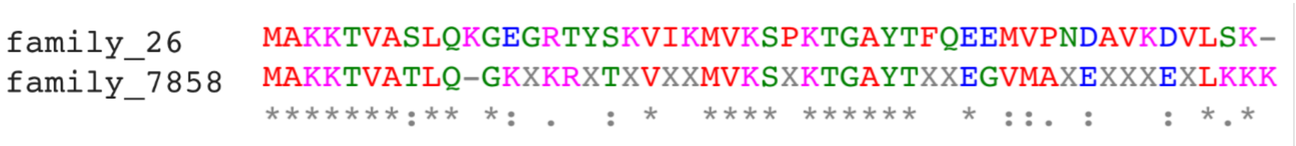


**Supplementary figure 5. Homology between family 26 and family 7858, potential novel ribosomal family of proteins.**

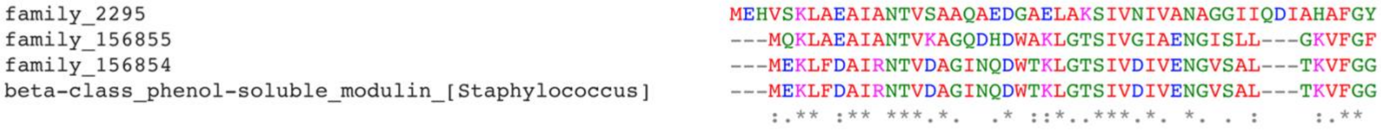


**Supplementary figure 6. Multiple sequence alignment between 3 families that are core to skin and a beta-class phenol-soluble modulin protein.**

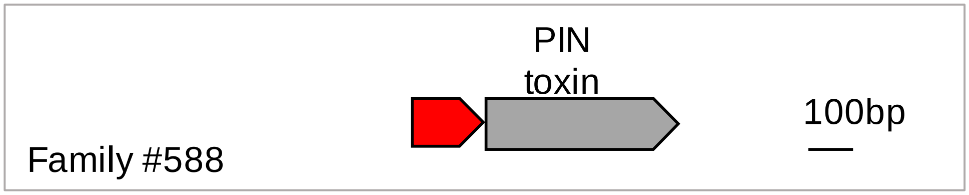


**Supplementary figure 7. Potential novel antitoxin small proteins.** Small proteins are predicted to be antitoxins of toxin-antitoxin systems based on their recurrent localization next to known toxin genes. This genomic organization was identified in 16/18 species in which we identified this small gene (Supplementary Table S5).

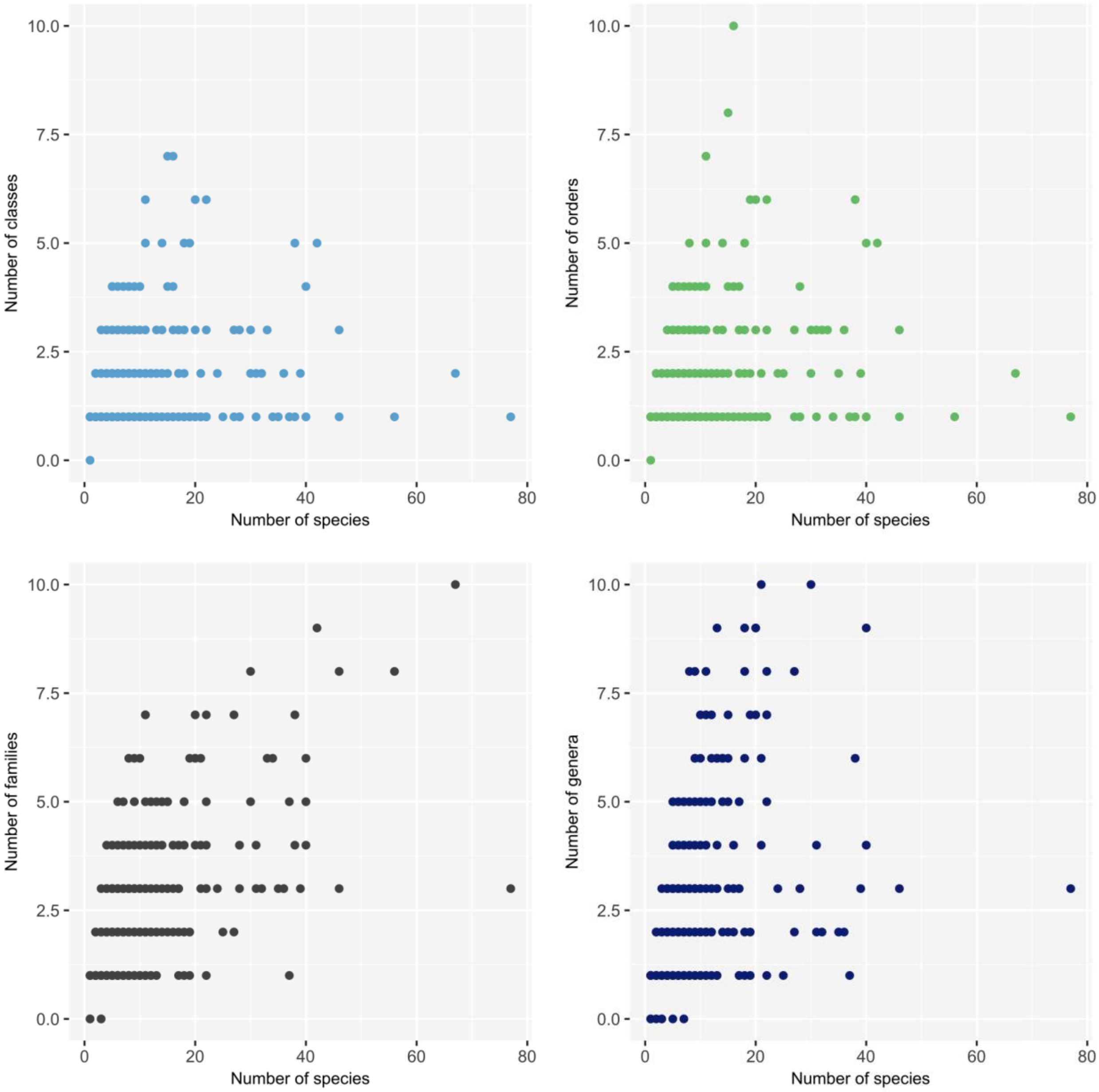

**Supplementary figure 8**. Scatter plots of 401 small protein families that are found in vicinity to HGT-associated genes. Each dot represents the number of species vs number of orders/classes/families/genera that encode for each family.

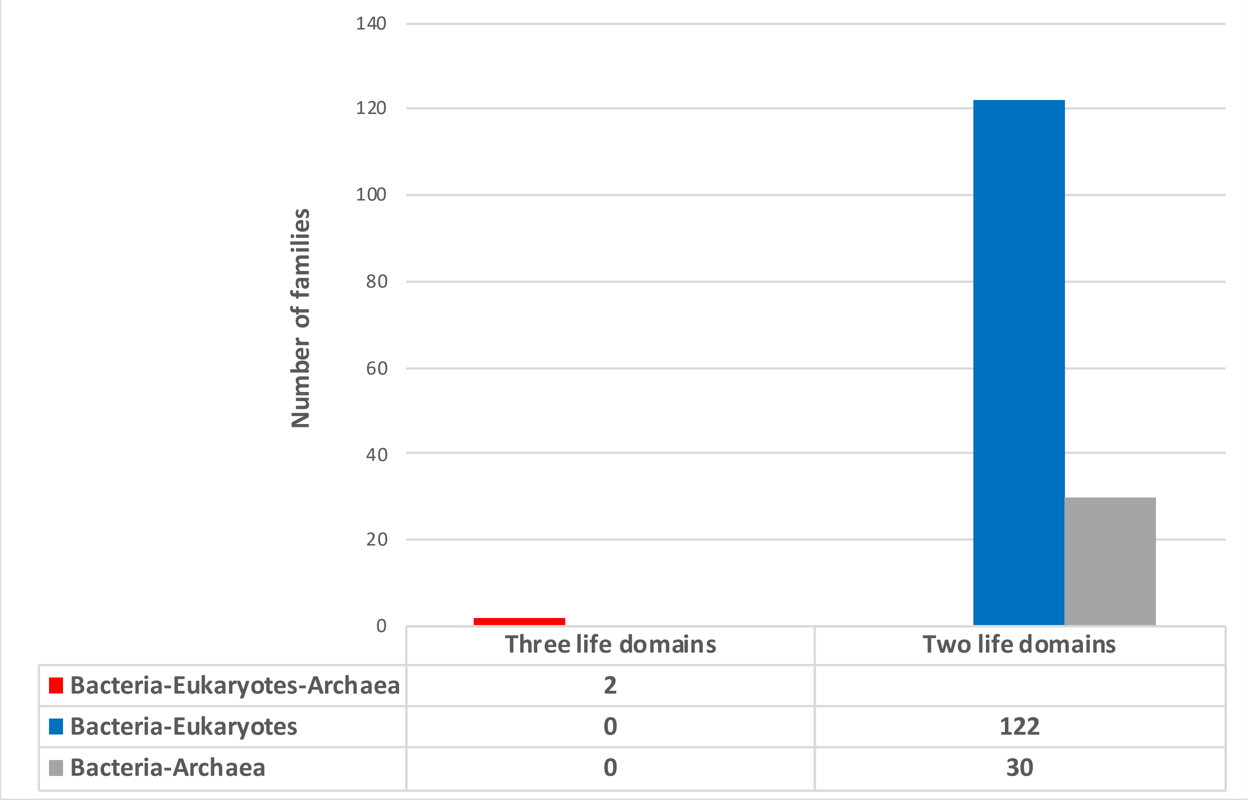


**Supplementary figure 9**. Numbers of small proteins families that were detected in multiple life domains.
